# Integrative Multi-Omics Identifies CDK1 as a Key Signaling Regulator of CD4^+^ T Cell Effector Function

**DOI:** 10.1101/2025.10.30.685551

**Authors:** Nila H. Servaas, Hanke Gwendolyn Bauersachs, Luisa Abreu, Annique Claringbould, Ivan Berest, Jennifer J. Schwarz, Frank Stein, Maria Fälth-Savitski, Lena Eismann, James P. Reddington, Judith B. Zaugg

## Abstract

CD4⁺ T cell differentiation is orchestrated by coordinated signaling, transcriptional, and epigenomic programs, yet how signaling connects to chromatin and genetic variation in human T cells remains unclear. Here, we generated an integrative multi-omics map of human CD4⁺ T cell activation and differentiation, combining phosphoproteomics, transcriptomics, and chromatin accessibility under Th0, Th1, and iTreg polarization. Within 10 minutes of activation, we observed rapid phosphorylation changes of RNA-binding proteins accompanied by degradation of effector-associated transcripts, preceding chromatin remodeling and later transcriptional activation of the same genes. Moreover, our data highlights how site-specific phosphorylation refines TF activity during T cell differentiation and activation, and identifies CDK1 as a regulator of Th1 effector function. Indeed, we found that a low dose of CKD1 inhibition impairs IFN-γ expression and pro-inflammatory differentiation, while preserving regulatory features in iTregs. Single-cell multi-omic profiling upon CDK1 inhibition revealed how CDK1 activity shapes subset-specific gene regulatory networks, which are enriched for genetic variants associated with immune-traits. Specifically, CDK1-sensitive TFs, including IRF8, connect immune trait heritability to enhancer accessibility at *IFNG* and *TNF* loci. Together, these results establish CDK1 as a signaling hub that couples phosphorylation to gene regulation and genetic risk, with therapeutic relevance in autoimmune disease.

## Introduction

T cells are central to immune homeostasis, with critical roles in anti-tumor immunity, defense against pathogens, and development of autoimmunity. CD4⁺ T helper (Th) cells differentiate into specialized subsets upon activation, including Th1, Th2, Th17, and regulatory T cells (Tregs)^1^. These subsets coordinate responses through cytokine secretion: Th1 cells produce interferon-γ (IFN-γ) to drive inflammation, while Tregs suppress activation to maintain tolerance^2^. Balanced differentiation is therefore essential for effective yet non-pathogenic immunity.

Naive CD4⁺ T cell differentiation is guided by extracellular cues that trigger intracellular signaling cascades and gene regulatory networks (GRNs)^3,4^. Lineage-defining transcription factors (TFs), including T-bet (Th1)^5^, and FOXP3 (Treg)^2^, reprogram the transcriptome and chromatin landscape. Disruption of these networks can contribute to immune dysregulation and impaired tumor immunity.

Recent single-cell transcriptomics, ATAC-seq, and CRISPR screens have mapped CD4⁺ T cell diversity and identified regulators of activation^3,6–8^. Yet, the upstream signaling events that initiate and sustain these programs remain poorly defined. Several signaling pathways are implicated in the differentiation of CD4⁺ T cells. mTORC1, for example, regulates Th1 differentiation by phosphorylating T-bet^9^, while IMPK promotes Treg differentiation^10^. However, an integrative map linking signaling cascades, TFs, and epigenomic remodeling in human CD4+ T cells is lacking. Moreover, although many immune-disease–associated single-nucleotide polymorphisms (SNPs) lie in noncoding regulatory regions active in T cells, how signaling pathways interface with genetic risk variants through chromatin and TF networks remains unclear. Previous phosphoproteomic studies have largely been restricted to mouse models^11–13^, or immortalized T cell lines^14,15^, while analyses of primary human T cells^16^ remain scarce.

To overcome these gaps we performed a time-resolved, integrative multi-omics analysis of primary human CD4⁺ T cell differentiation in healthy donors. Naive cells were polarized into Th1 and induced regulatory T cells (iTregs), followed by profiling of the transcriptome (RNA-seq), chromatin accessibility (ATAC-seq), and phosphoproteome (mass spectrometry) across activation and differentiation time points to map signaling and gene regulatory programs. Kinase enrichment analyses identified cyclin-dependent kinase 1 (CDK1) as a previously unrecognized regulator of Th1 effector function. Functional perturbation with a selective CDK1 inhibitor, coupled with single-cell multi-omic profiling, demonstrated that CDK1 activity is required for pro-inflammatory effector function while sparing regulatory programs. Finally, by intersecting TF-associated chromatin accessibility with genome-wide association study (GWAS) data, we show that CDK1-sensitive regulatory elements are enriched for immune trait heritability, linking phosphorylation dynamics to transcriptional control and genetic risk.

Our findings reveal a signaling–epigenetic–genetic axis in human T cells and highlight the power of integrative multi-omics to uncover regulators of immune fate decisions and connect them to complex disease genetics.

## Results

### Multi-omics time-course profiling of CD4⁺ T cell activation and differentiation

To dissect the signaling and gene regulatory events underlying CD4⁺ T cell activation and differentiation, we generated an *in vitro* differentiation time-course using naive CD4⁺ T cells from three healthy human donors (**Figure 1A**). Cells were polarized to Th0 (anti-CD3/CD28, IL-2), Th1 (anti-CD3/CD28, IL-2, IL-12, anti-IL-4), and iTreg (anti-CD3/CD28, IL-2, TGF-β) and cultured for five days to allow lineage commitment. Intracellular flow cytometry confirmed robust IFN-γ induction in Th1 cells and FOXP3 in iTregs (**Figure 1B**).

**Figure 1.**
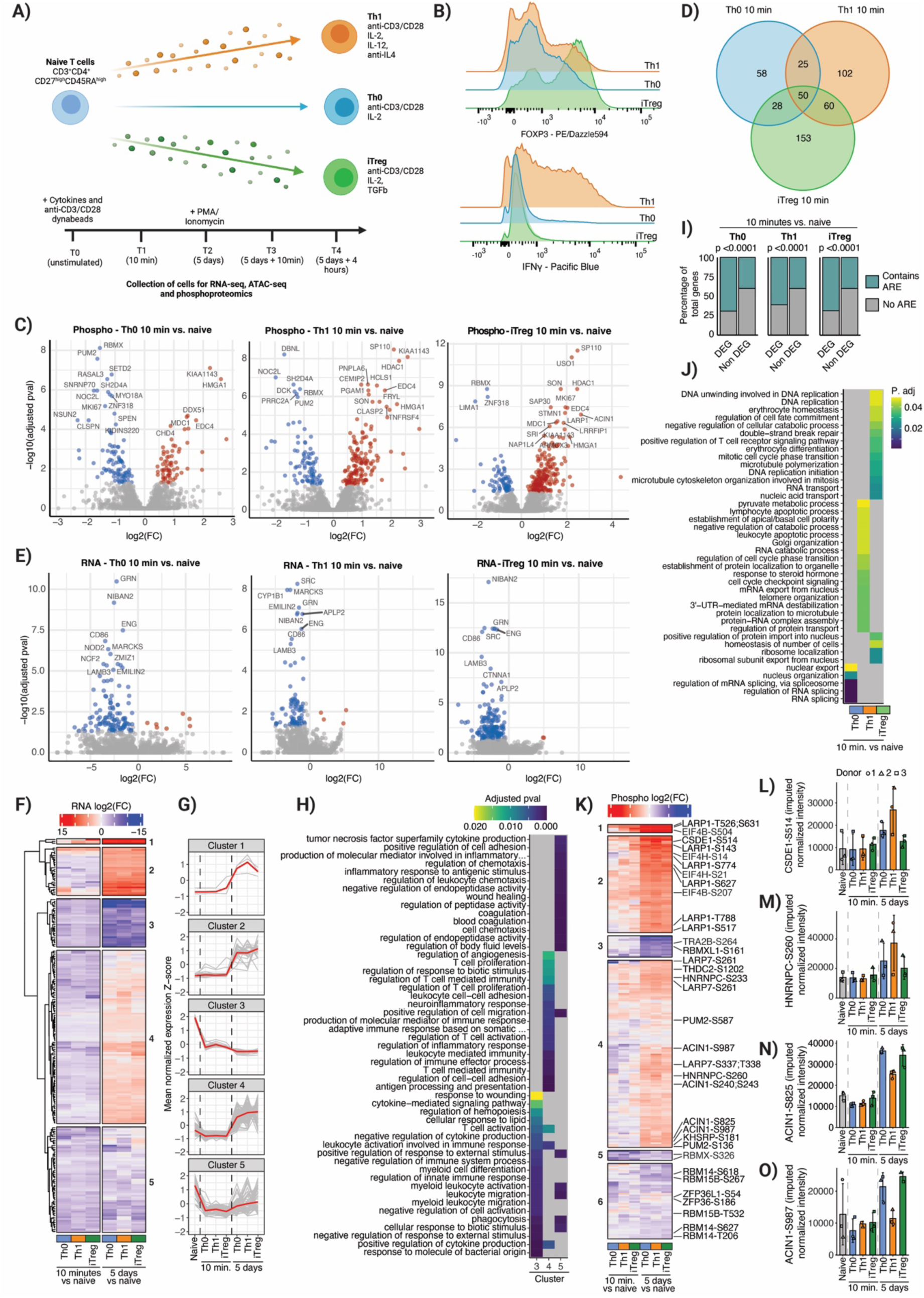
Integrated phosphoproteomic and transcriptomic profiling reveals layered post-transcriptional regulation in early T cell activation. (**A**) Experimental design: Naive CD4⁺ T cells were stimulated and polarized to Th0, Th1, or iTreg, and profiled at the indicated time points by phosphoproteomics, RNA-seq, and ATAC-seq. (**B**) Flow cytometric validation of subset identity (FOXP3, IFN-γ) at day 5 after 4h PMA/ionomycin restimulation. (**C**) Volcano plots of differentially phosphorylated peptides (10 min vs. naive) in Th0, Th1, and iTreg (red: up, blue: down; adj. p < 0.05). (**D**) Venn diagram of significantly regulated phosphosites (10 min vs. naive) across subsets. (**E**) Volcano plots of RNA-seq (10 min vs. naive) per subset. (**F**) Clustering of differentially expressed transcripts across 10 min and 5 days into five temporal patterns. (**G**) Line plots showing normalized expression (Z-score) of individual genes (grey lines), and the cluster mean (red lines). (**H**) GO term enrichment for gene clusters identified in (F). (**I**) Proportion of downregulated transcripts containing AU-rich elements (AREs) compared to non-differential transcripts. (**J**) Enriched GO terms among early upregulated phosphosites (10 min vs. naive). Top 15 terms (based on adj. p) for each comparison are shown. (**K**) Heatmap of differentially phosphorylated RNA-binding proteins (RBPs) across 10 minutes and 5 days. Six clusters reflect distinct temporal modules. (**L–O**) Bar plots of phosphorylation intensity (mean ± SD) for selected RBPs with individual donors shown.

We profiled five key time points: (1) naive baseline, (2) 10 minutes after initial stimulation, (3) five days after differentiation, and five-day differentiated cells restimulated with PMA (phorbol 12-myristate 13-acetate) and ionomycin for (4) ten minutes or (5) four hours. At each time point, we profiled bulk RNA-seq, ATAC-seq, and phosphoproteomics to compare acute signaling during initial activation against secondary stimulation in differentiated cells.

We detected 4,445 unique phosphopeptides, of which 2,378 (on 1,174 proteins) were reproducibly observed across at least two donors and retained for analysis. Principal component analysis (PCA) of the top 500 variable phosphosites (**Supplemental Figure 1A**) revealed a major shift along PC1 (44.6%), separating naive and early activated cells (0 and 10 min) from differentiated states (5 days+), thus reflecting extensive remodeling during differentiation. PC2 (9.4%) captured T cell subset- and time-dependent diversity. Similar temporal and subset-specific separation was observed for RNA-seq (27,930 genes; PC1 69.5%, PC2 10.7%; **Supplemental Figure 1B**) and ATAC-seq (110,842 peaks; PC1 67.3%, PC2 11.7%; **Supplemental Figure 1C**).

### Phosphorylation of RNA-binding proteins drives early decrease in RNA levels linked to CD4⁺ T cell activation

Ten minutes after stimulation, we observed widespread phosphoproteomic remodeling compared to naive CD4⁺ T cells (**Figure 1C**), reflecting immediate phosphorylation events triggered by T cell receptor (TCR) and cytokine signaling. Of the 476 differentially phosphorylated sites, 164 (34.2%) were significant in all three subsets. While additional sites appeared subset-specific by statistical cutoffs (p. adj < 0.05, **Figure 1D**), fold-changes were highly correlated between Th0, Th1, and iTregs (**Supplemental Figure 1D**), indicating that the earliest responses largely reflect shared TCR signaling rather than subset-specific programs. At this stage, chromatin accessibility remained unchanged, as expected given the slower kinetics of epigenomic remodeling. In contrast, we observed significant downregulation of multiple RNA transcripts (**Figure 1E**), which were enriched in pathways related to regulation of T cell activation (**Supplemental Figure 1E**). Notably, this mRNA downregulation was absent upon 10 minutes of restimulation with PMA/ionomycin at day 5, suggesting that early transcript suppression is specific to primary activation.

To assess how these early changes evolved, we clustered genes differentially expressed at 10 minutes after initial stimulation and tracked their expression at 5 days. This revealed five major temporal expression patterns, three of which (cluster 1, 3 and 4) were characterized by an initial downregulation at 10 minutes (**Figure 1F/G**). Genes in the largest cluster (cluster 4) were suppressed at 10 minutes but exceeded baseline expression after 5 days, and were enriched in antigen presentation, T cell–mediated immunity, and proliferation, suggesting a delay before full effector engagement. Supporting the notion that early signaling primes later effector engagement, we observed phosphorylation of signaling regulators, including CARD11 (S593, required for NF-κB activation downstream of the TCR^17^) and PTPN7 (S143, a modulator of MAPK^18^) (**Supplemental Figure 1F**). Cluster 3 showed early repression after 10 minutes that strengthened after 5 days, enriched for genes implicated in negative regulation of cytokine production and T cell activation, consistent with silencing of feedback regulators during differentiation. Cluster 5 remained repressed and was enriched for inflammatory signaling, chemotaxis, and TNF production, consistent with tight post-transcriptional regulation of pro-inflammatory programs in T cells^19^. Together, these clusters highlight that early transcript suppression follows distinct trajectories, with some pathways recovering and others remaining repressed during differentiation.

To investigate the mechanisms underlying the early transcript downregulation observed at 10 minutes, we examined sequence features of the affected genes. They were significantly enriched for adenylate-uridylate-rich elements (AU-rich elements; AREs; annotations from AREsite2^20^; **Figure 1I**), which are canonical targets of RNA-binding proteins (RBPs) that control splicing, translation, mRNA stability and degradation^21^. Consistently, early differentially phosphorylated proteins were enriched for RNA metabolism pathways, including splicing, transport and 3’-UTR-mediated decay (**Figure 1J**).

To further explore the role of RBPs, we curated a list of RBPs from the RNA-Binding Protein DataBase (RBPDB)^22^ and focused on those with significant differential phosphorylation in any pairwise comparison. Clustering RBPs based on phosphorylation changes at 10 minutes and 5 days relative to naive cells (the timepoints showing the strongest shifts, **Supplemental Figure 1G**), revealed six distinct clusters (**Figure 1K**). RBP clusters 1 and 2 (early and sustained activation) contained RBPs highly phosphorylated at 10 minutes and maintained at 5 days, consistent with sustained post-transcriptional regulation. These included LARP1 (an mTORC1 target regulating mRNA translation^23,24^), and CSDE1 (modulates immune transcript stability and translation^25^), which showed strongest phosphorylation in Th1 cells at day 5 (**Figure 1L**). EIF4B/H (core mTOR-regulated translation factors^26^), were also present, underscoring rapid engagement of translational control that persists into differentiation. RBP cluster 3 (late repression) comprised RBPs with reduced phosphorylation at day 5, including splicing factors TRA2B, (regulator of TCR sensitivity^27^), and RBMXL1^28^, suggesting reduced alternative splicing in differentiated cells. RBP cluster 4 (delayed activation) showed increased phosphorylation at day 5, reflecting a second wave of post-transcriptional control that may refine effector programs. Members included LARP7 (RNA Pol II elongation^29^), KHSRP^30^ and PUM2^31^ (mRNA decay) and m6A-associated factors YTHDC2^32^ and HNRNPC^33^. Notably, HNRNPC phosphorylation at S260 was highest in Th1, suggesting subset-specific splicing to support inflammatory gene expression (**Figure 1M**). ACIN1 (a splicing factor linked to cell cycle control^34^), was phosphorylated in Th0 and iTregs but less in Th1 (**Figure 1N/O**), further highlighting subset-specific regulation. RBP clusters 5 and 6 (early and progressive repression) included splicing factor RBMX^28^, m6A methyltransferase complex member RBM15B^35^, and ZFP36/ZFP36L1 (regulators of immune transcript decay in T cells^19,36^). Their dephosphorylation likely reflects reduced RNA-binding-mediated decay and translational repression, thereby stabilizing effector transcripts as differentiation proceeds. Reduced phosphorylation of RBM14 and THUMPD1 (involved in RNA processing and modification^37,38^), further supports a transition toward a more stable transcriptome in differentiated cells.

Together, these patterns highlight dynamic phosphorylation of RBPs as a key layer of post-transcriptional regulation that fine-tunes CD4⁺ T cell differentiation.

### Post-translational and chromatin-based regulation shape TF activity during CD4⁺ T cell differentiation

After five days of polarization under Th0, Th1, or iTreg conditions, CD4⁺ T cells were committed to distinct effector phenotypes (**Figure 1B**), enabling dissection of subset-specific regulatory programs. Compared to naive cells, we observed extensive transcriptional remodelling (**Supplemental Figure 2A**), and widespread chromatin accessibility changes (**Supplemental Figure 2B**). Promoter accessibility and gene expression changes correlated strongly at day 5 and after 4 hours of restimulation (**Figure 2A**), indicating that TFs drive both differentiation-specific and secondary activation responses. Preceding the chromatin remodeling, at 10 minutes post restimulation we observed phosphorylation of proteins linked to chromosome organization, spindle dynamics, and chromatin remodeling (**Supplemental Figure 2C**), including CHD1 (S1677) and SMARCA4 (S1417) (**Supplemental Figure 2D**), pointing to rapid signaling inputs that may prime later epigenomic responses. In contrast, minimal changes in RNA or chromatin were observed at this early restimulation time point. This points to a layered regulatory architecture, with RBPs shaping immediate transcript stability, followed by TF- and chromatin remodeling that regulate subset-specific programs.

**Figure 2.**
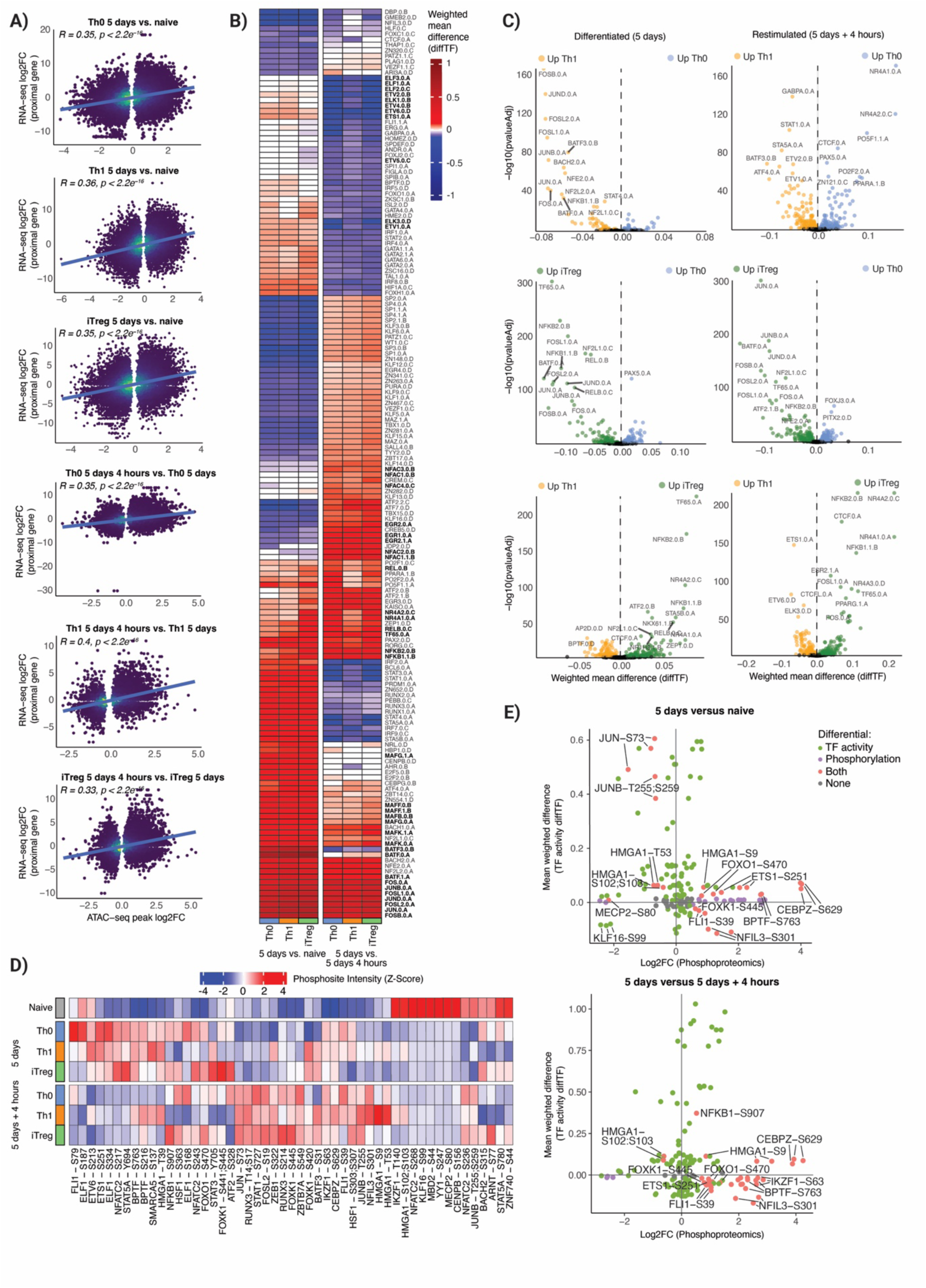
Integrated analysis of TF activity, gene expression, and phosphorylation reveals temporal regulation during Th subset differentiation and restimulation. **(A)** Scatter plots showing the correlation between chromatin accessibility (ATAC-seq peak log2FC) and gene expression (RNA-seq log2FC) across conditions. Blue regression lines and Pearson’s correlation coefficients (R) indicate correlation between accessibility and expression across conditions. **(B)** Heatmap showing the top 100 TFs ranked by absolute DiffTF activity differences for each comparison. Left panel: TF activity differences between day 5 and naive (T0). Right panel: TF activity differences between 5 days and 5 days + 4 hours. Color intensity indicates weighted mean difference as estimated by diffTF for top 50 induced and top 50 reduced TFs. AP-1, NF-κB, EGR, ETV, NFAT and NR4A family members are highlighted in bold. **(C)** Volcano plots of TF activity differences (diffTF) for selected pairwise comparisons at 5 days (left) and 5 days + 4 hours (right). TFs significantly upregulated in Th0, Th1, or iTreg subsets are colored accordingly. Vertical dashed lines mark zero activity difference. **(D)** Heatmap of phosphoproteomics data showing the Z-score-normalized intensity of selected phosphosites across subsets and time points. Conditions include naive (T0), differentiated cells at day 5, and restimulated cells at day 5 + 4 hours. Phosphorylation changes are visualized for all TFs detected according to the HOCOMOCO database. **(E)** Scatter plots showing the integration of TF activity (diffTF) and phosphosite-level regulation. Top: Day 5 vs. naive; bottom: Day 5 + 4h vs. day 5. TFs showing both phosphorylation and chromatin activity changes are highlighted in red.

To further study TF dynamics, we inferred differential TF activity using diffTF^39^. AP-1 (FOS/JUN/JUND/BATF/MAFs) and NF-κB family members (NFKB1/2, TF65, REL) were strongly induced during differentiation and restimulation, consistent with their roles in TCR-driven chromatin remodeling^40,41^ (**Figure 2B**). Immediate-early factors EGR1/2^42^ decreased at five days, reflecting a shift from transient activation to stable commitment. ETS family member activity (ETS/ETV/ELF/ELK) decreased following restimulation, in line with their repressive roles in immune differentiation^43,44^. Restimulation prominently induced NFAT (NFAC1–4) and NR4A1/2, factors that drive effector gene expression and enforce feedback upon TCR signalling^45,46^.

We next compared TF activity across subsets (**Figure 2C**). Both Th1 and iTreg showed stronger AP-1 and NF-κB activity than Th0, consistent with engagement of robust activation programs in differentiated states. NF-κB activity was particularly elevated in iTregs, in line with its role in Treg stability and suppressive function^47^, and was accompanied by increased activity of known Treg-associated TFs including STAT5B^48^, NR4A1/2^49^, and PPARG^50^. iTregs also displayed higher activity of ZEP1 and FOSL1, highlighting potential novel regulators. ETS1, a known Th1 regulator^51^, was more active in Th1 cells, as well as AP2D and ELK3, pointing to novel candidate TFs driving Th1 differentiation.

Because TF activity is often post-translationally regulated^52^, we examined TF phosphorylation dynamics across naive, differentiated (5 days), and restimulated (5 days + 4h) cells, identifying 51 differentially phosphorylated sites (**Figure 2D**). ETV family members showed reduced phosphorylation after restimulation, consistent with their decreased activity. Notably, we detected phosphorylation of ETS1 at S251, which reinforces its autoinhibition and limits DNA binding^53^ (**Supplemental Figure 2E**). Several TFs showed subset-specific regulation: STAT5A and BACH2 (critical for Tregs^54,55^) were reduced in Th1 cells but maintained or elevated in Th0 and iTregs, alongside ZNF740 and ARNT. Conversely, HMGA1 phosphorylation increased at multiple sites specifically in Th1, consistent with its reported role in *IFNG* regulation^56^.

To directly link phosphosite dynamics with function, we compared phosphorylation with motif accessibility–inferred activity (**Figure 2E**). JUNB phosphorylation at T255/S259 (a degradation motif^57^), decreased as JUNB activity increased (**Supplemental Figure 2F**), suggesting stabilization during differentiation. HMGA1 activity rose in parallel with reduced phosphorylation at S102/S103, sites known to impair DNA binding^58^ (**Supplemental Figure 2G**). NFKB1 activity correlated with increased phosphorylation at S907, a site fine-tuning NF-κB signaling^59^, with both signals strongest in iTregs (**Supplemental Figure 2H)**. These cases illustrate how site-specific phosphorylation can act as a regulatory switch, refining TF activity during T cell differentiation and activation.

### Network-based phosphoproteomic analysis reveals candidate kinases shaping CD4⁺ T cell activation and differentiation

To identify kinases driving phosphorylation changes during CD4⁺ T cell differentiation and reactivation, we applied kinase set enrichment analysis (KSEA)^60^, which relies on known kinase-substrate relations, to our dataset (**Figure 3A**). Cyclin-dependent kinases (CDKs) were strongly enriched in differentiated and restimulated cells, reflecting their roles in proliferation. Strikingly, CDK1 showed the highest inferred activity specifically in Th1 cells at day 5 and after restimulation (**Supplemental Figure 3**), suggesting a role in supporting Th1 effector programs. We also observed increased activity of kinases important in T cell signalling, including CSNK2A1^61^, PRKCA/D^62^, AKT1^63^, and mTOR^24,64^.

**Figure 3.**
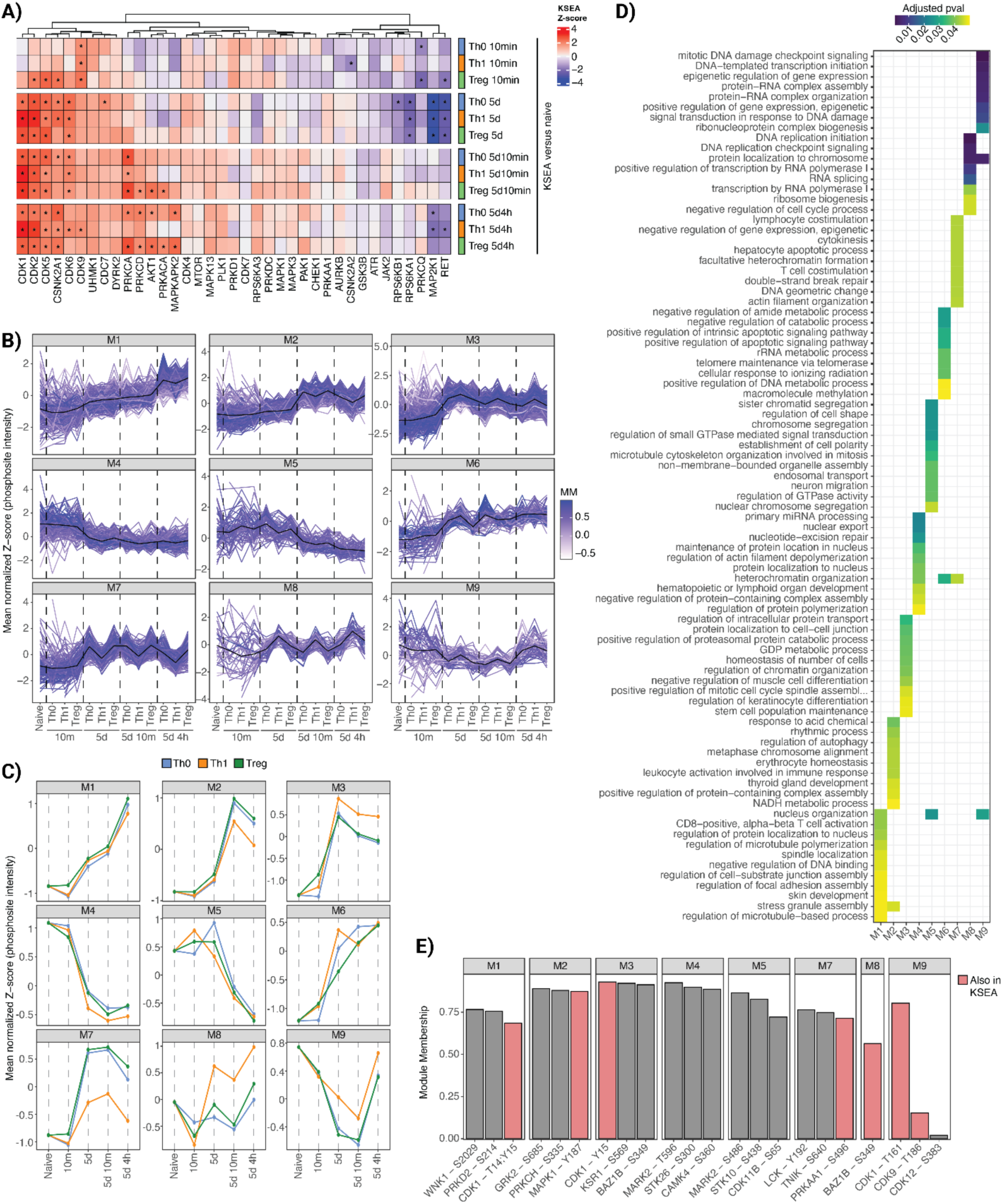
Identification of dynamic signaling modules and their candidate regulators during T cell differentiation and restimulation. (**A**) Kinase activity scores inferred by KSEA (kinase–substrate enrichment analysis) across time points and T cell subsets. Each column represents a kinase, and color scale reflects relative Z-scores of predicted activity versus naive. Asterisks indicate significant activation (NetworKIN score cutoff ≥ 5, adjusted p-value < 0.05, and minimum of 3 substrates). Subset and timepoint are indicated in the annotation bars. (**B**) Temporal profiles of phosphosite modules (M1–M9) derived from WGCNA across all conditions. Each line represents a phosphosite, colored by its module membership (MM).

Complementary to KSEA, we employed weighted gene co-expression network analysis (WGCNA)^65^, applied directly to phosphosite-level data, which identified nine modules of co-regulated phosphosites, representing signaling programs independent of prior knowledge (**Figure 3B**). Module-level phosphorylation dynamics (**Figure 3C**), revealed temporal and T cell subset-specific patterns. Activation-associated modules (M1–M3, M6, M7) increased in differentiated and restimulated cells and were enriched for processes including stress granule assembly and CD8⁺ T cell activation (M1), leukocyte activation (M2), chromatin organization (M3), and cytokinesis and lymphocyte costimulation (M7) (**Figure 3D**). Modules linked to RNA metabolism (M8, M9) were strongly dephosphorylated during initial stimulation and regained phosphorylation during differentiation and restimulation, especially in Th1 cells. These dynamics align with our observations of RBP phosphorylation and ARE-transcript regulation (**Figure 1F–J**), suggesting that post-transcriptional control is tightly coupled to effector function.

Kinase module membership scores, which measure how closely kinase phosphorylation patterns align with overall module dynamics, identified CDK1 among the top-3 kinases in M1 and M3 (**Figure 3E**), associated with activation and chromatin organization, as well as the top kinase in M9, enriched for RNA-related processes. Both M3 and M9 showed elevated phosphorylation in Th1 cells, reinforcing the hypothesis that CDK1 contributes to Th1-specific signaling. This aligns with our KSEA results, where CDK1 showed the strongest enrichment in Th1 cells compared to Th0 and iTregs (**Figure 3A**).

Dashed vertical lines indicate stimulation time points. T cell subsets are displayed sequentially on the x-axis for visualization but do not represent a developmental progression. (**C**) Mean normalized phosphorylation intensity (Z-score) for each module across naive, differentiated (5 days), and restimulated (5 days + 4 hours) Th0, Th1, and iTreg cells. (**D**) GO enrichment analysis of phosphoproteins within each module. (**E**) Candidate kinases associated with each phospho-module (M1–M9), based on module membership scores. Highlighted in red are kinases that also showed significant enrichment in KSEA analysis (A).

### CDK1 is implicated in Th1 effector function independently of its role in cell cycle

Given its consistent enrichment across KSEA and WGCNA analyses, and strong association with Th1-related phospho-modules, we tested the functional role of CDK1 during CD4⁺ T cell differentiation. Naive CD4+ T cells were polarized in the presence or absence of RO-3306, a selective CDK1 inhibitor^66^ (**Figure 4A**). To avoid confounding effects from CDK1’s canonical cell cycle role, we performed a dose–response titration (0.15–10 μM) and observed that concentrations ≤2.5 μM preserved viability and proliferation **(Figure 4B–C**).

**Figure 4.**
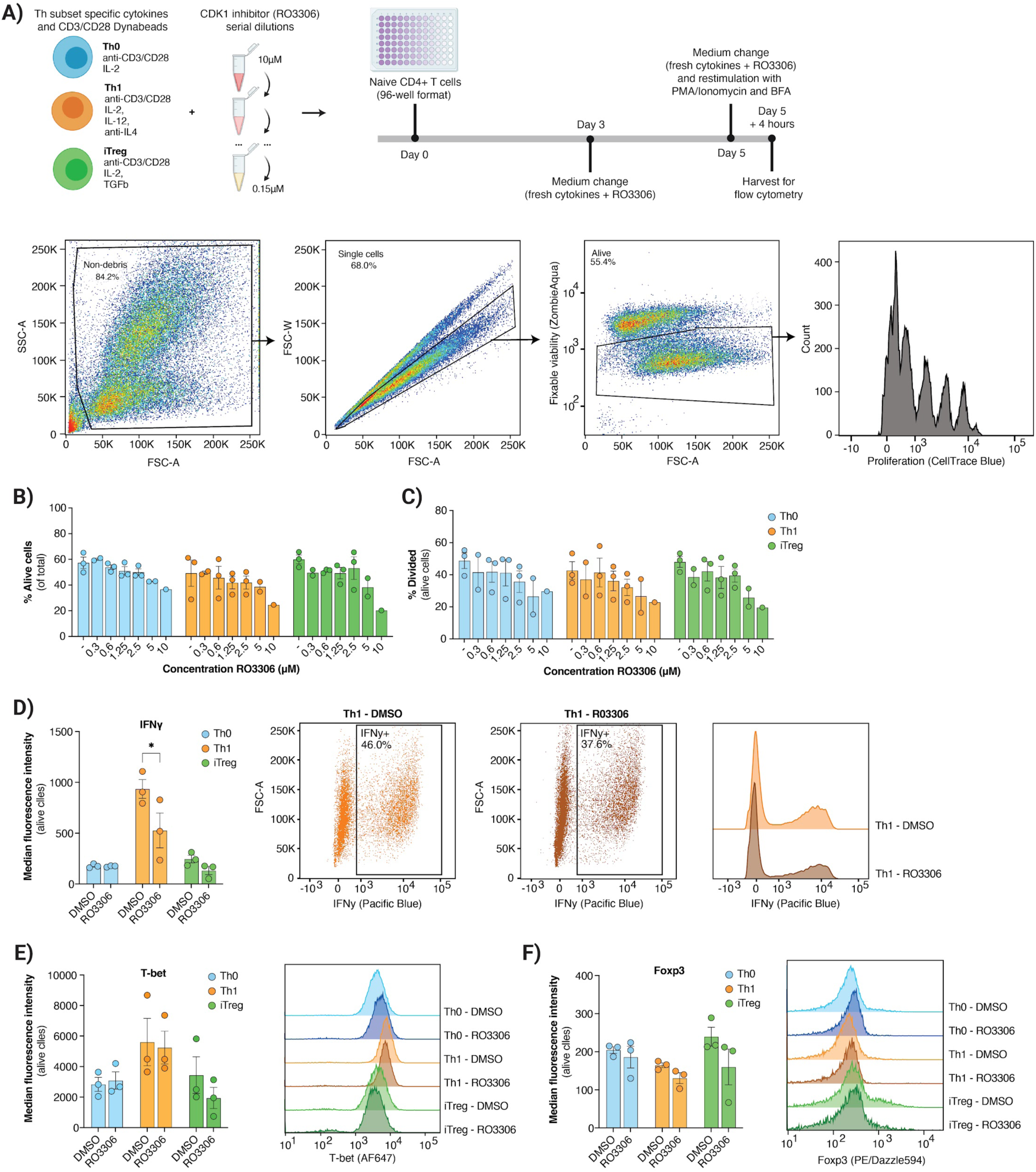
CDK1 regulates Th1 effector function independently of cell cycle. **A)** Experimental design for CDK1 inhibition assays. Naive CD4⁺ T cells were stimulated under Th0, Th1, or iTreg polarizing conditions with serial dilutions of CDK1 inhibitor RO3306 (0.15– 10 μM). Cells were cultured for 5 days with a media change on day 3, followed by 4 hour restimulation with PMA/ionomycin and brefeldin A (BFA) on day 5. **(B)** Percentage of live cells (flow cytometry) after 5 days of culture with RO3306, across Th0 (blue), Th1 (orange), and iTreg (green) conditions. Dots represent individual experiments with cells from different donors. (**C**) Percentage of divided cells (among live cells) across conditions, measured by CellTrace™ Blue staining. (**D**) Intracellular expression IFN-γ, (**E**) T-bet and FOXP3 (**F**) following CDK1 inhibition. Bar plots show median fluorescence intensity (MFI) in each subset (*p < 0.05).

We next assessed how CDK1 inhibition affects the expression of canonical Th1 and Treg markers using intracellular flow cytometry after 5 days of differentiation followed by 4 hours restimulation with PMA/ionomycin. In line with our phosphoproteomic predictions, CDK1 inhibition led to a significant reduction of IFN-γ in Th1 cells (**Figure 4D**). Expression of the canonical Th1 TF T-bet was unchanged (**Figure 4E**), indicating CDK1 does not block Th1 lineage commitment but impairs effector function. FOXP3 expression in iTregs was also not significantly affected (**Figure 4F**). These results support a role for CDK1 in modulating Th1 function, beyond its canonical role in cell cycle.

### Single-cell multi-omics reveals divergent effects of CDK1 inhibition on effector and regulatory T cell programs

To dissect how CDK1 shapes CD4⁺ T cell differentiation and activation, we generated joint scRNA-seq and scATAC-seq data using Single-cell Ultra-high-throughput Multiplexed sequencing (SUM-seq)^8^, across three differentiation conditions (Th0, Th1, iTreg), two time points (day 5 and day 5 + 4h restimulation), and three CDK1 inhibitor regimens (DMSO control, continuous 2.5 µM RO-3306, and 2.5 µM RO-3306 only during final activation) in naive CD4⁺ T cells from five healthy donors. After 5 days of differentiation, cells were left unstimulated or restimulated with PMA/ionomycin for 4 hours. To disentangle effects on differentiation versus activation, we compared continuous inhibition throughout culture and restimulation (“RO-3306 full”) with inhibition restricted to the final activation (“RO-3306 late”) (**Figure 5A**).

**Figure 5.**
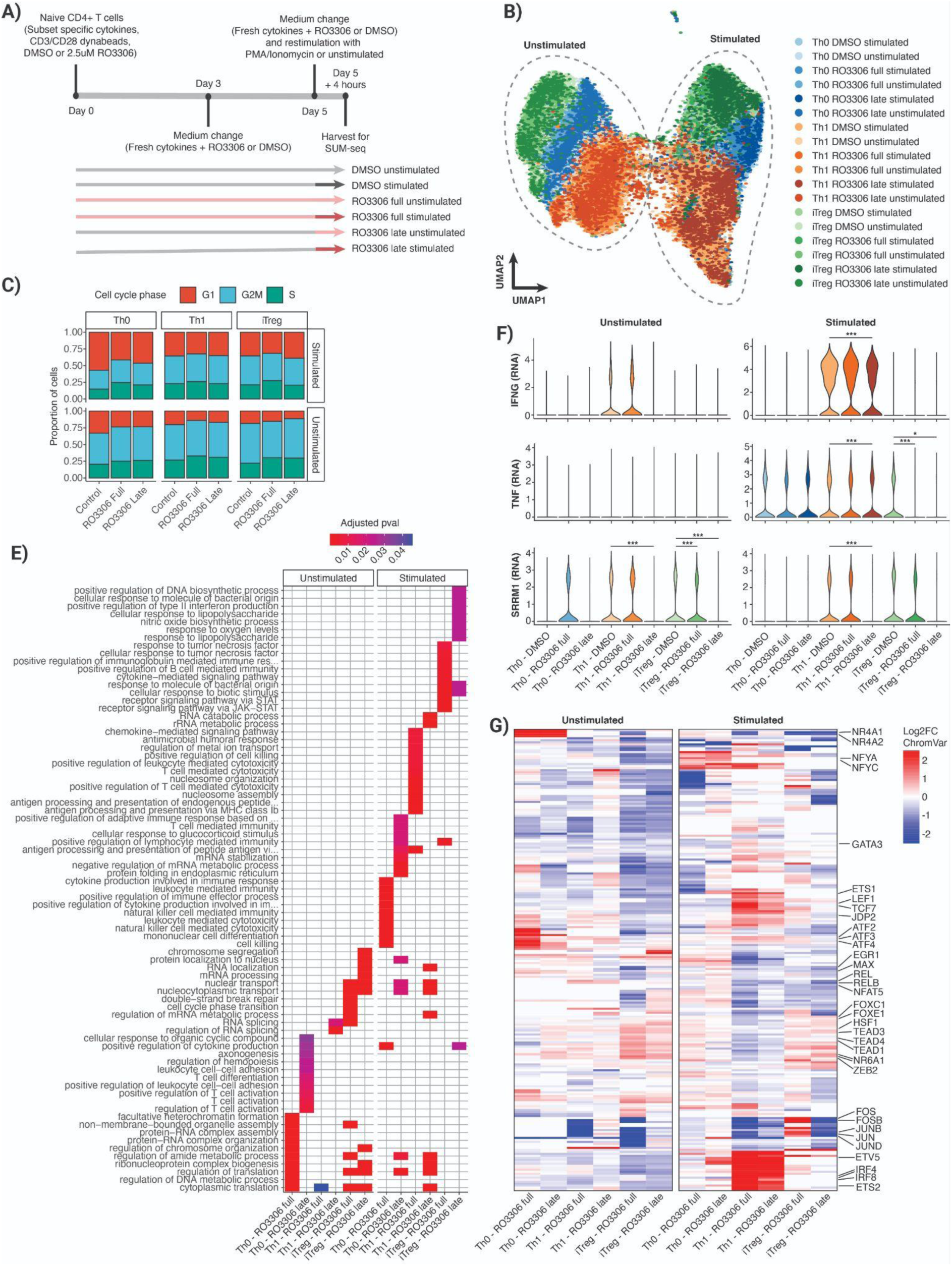
CDK1 inhibition reshapes transcriptional response in human CD4⁺ T cells. **(A)** Schematic overview of the experimental setup. Naive CD4⁺ T cells were differentiated into Th0, Th1, and iTreg subsets in the presence of CDK1 inhibitor RO-3306 or DMSO control, either continuously (“Full”) or only during the final 4h (“Late”), followed by restimulation or no restimulation (4 hours PMA/ionomycin) prior to SUM-seq profiling. **(B)** Weighted nearest neighbours UMAP of integrated scRNA- and scATAC-seq data showing all cells colored by condition. **(C)** Stacked bar plots showing the distribution of cells across cell cycle phases (G1, S, G2/M) in each condition and T cell subset. **(E)** GO term enrichment of differentially expressed genes in the different CDK1 inhibition conditions versus DMSO control. For each condition, the top 10 terms (based on adjusted p value) are shown. **(F)** Violin plots of scRNA-seq data for selected genes demonstrate in stimulated and unstimulated cells upon RO-3306 treatment. **(G)** Heatmap of differential TF activity (based on chromVAR scores) for CDK1 inhibition conditions versus DMSO control. Selected TFs are highlighted.

After quality control and Harmony integration, we retained 58,346 cells with paired RNA and ATAC profiles across 87 samples. UMAP visualization separated cells by subset and stimulation state (**Figure 5B**), with donors and replicates well mixed (**Supplemental Figure 4A/B**).

We first assessed cell cycle states based on canonical RNA signatures. Although CDK1 inhibition might be expected to induce G2/M arrest, treated cells showed no significant differences compared to controls, indicating that the selected dose did not impair cell cycle (**Figure 5C**). Restimulation with PMA/ionomycin induced a general shift toward G1 and away from G2/M across all subsets (**Supplemental Figure 4C)**.

Transcriptomic analysis revealed that CDK1 inhibition altered hundreds of genes across subsets, with substantial overlap across conditions (**Supplemental Figure 4D**). GO-enrichment highlighted pathways in T cell activation, cytokine and JAK–STAT signaling, and RNA metabolism (**Figure 5E**), suggesting that CDK1 regulates both immune programs and post-transcriptional processes. Consistent with flow cytometry (**Figure 4D**), *IFNG* was significantly reduced in stimulated Th1 cells, particularly under late inhibition, and *TNF* was broadly downregulated in Th1 and iTregs (**Figure 5F**)*. SRRM1*, an RBP linked to JAK–STAT signaling^67^, was also decreased (**Figure 5F**), reinforcing CDK1’s role in RNA processing.

We next inferred TF activity from scATAC data using chromVAR (**Figure 5G, Supplemental Figure 4E**). CDK1 inhibition broadly reduced AP-1 family TFs (FOS, FOSB, JUN, JUNB/D), consistent with impaired T cell activation. Indeed, CDK1 phosphorylates JUNB during mitosis^68^. Continuous inhibition paradoxically increased AP-1 activity in iTregs, consistent with context-dependent roles of AP-1 in promoting FOXP3 expression^69^. NF-κB factors (REL, RELB) also showed reduced activity in stimulated Th1s and iTregs, mirroring *TNF* downregulation (**Figure 5F**) and consistent with the central role of NF-κB in inflammatory programs^70^. Although NF-κB is not a known CDK1 substrate, CDK1 can indirectly modulate this pathway through adaptor phosphorylation^71^.

In parallel, CDK1 inhibition enhanced TFs associated with regulatory or alternative fates, including, including TEAD1/3/4^72^, HSF1^42^, and ZEB2^73^ in Th0 and iTregs (**Figure 5G**), supporting maintenance of a regulatory phenotype. In Th1s, inhibition preserved activity of lineage TFs such as ETV5 and ETS1, (also implicated in Treg biology^74^), but also activated competing programs. These included IRF4 and IRF8 (balancing effector and regulatory circuits^75–77^), LEF1 and TCF7 (driving T follicular helper differentiation^78^), and GATA3 (a Th2 regulator antagonizing Th1 identity^79^). Thus, CDK1 is not required for Th1 lineage specification but is essential for reinforcing the effector program: without it, Th1 cells retain partial identity yet increasingly engage alternative transcriptional circuits, coinciding with reduced IFN-γ expression and diminished AP-1 activity.

### Integration of gene regulatory networks and GWAS highlights CDK1-sensitive TFs at immune risk loci

To further characterize the roles of the CDK1-sensitive TFs we inferred an enhancer-driven GRN using SCENIC+^80^ to study their target genes across all subsets, stimulation states, and inhibitor conditions. The GRN captured both differentiation- and activation-associated programs, revealing enhancer-mediated regulation sensitive to CDK1 inhibition (**Figure 6A**).

**Figure 6.**
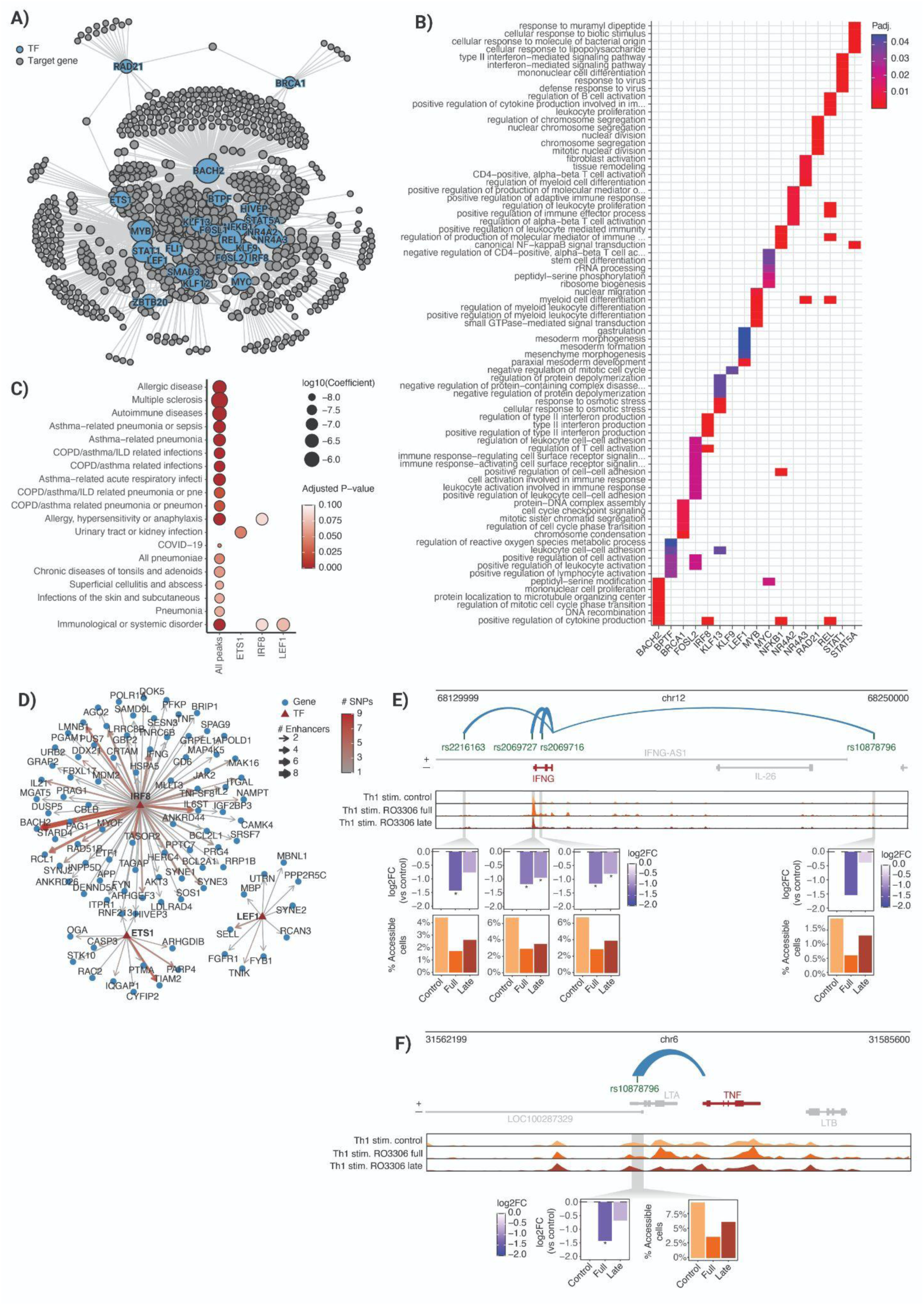
CDK1-sensitive TFs connect enhancer regulation to immune trait heritability. **(A)** Genome-wide TF–target GRN inferred from integrated scRNA– and scATAC-seq data. Blue nodes: TFs; grey nodes: target genes; edges represent enhancer-mediated links. **(B)** Gene Ontology enrichment of GRN target genes. Top 10 terms per TF are shown, colored by adjusted P-value. (**C**) Stratified LD score regression (S-LDSC) showing heritability enrichment of immune traits in TF-regulated peaks. Dot size indicates log₁₀-transformed effect size; color reflects FDR-adjusted P-value. (**D**) Subnetwork for IRF8, ETS1, and LEF1 linking disease-associated SNPs (red gradient) to enhancers (edges) and downstream genes (grey). Edge thickness indicates number of enhancers; edge color shows total SNP count. (**E/F**) Example loci where enhancers harboring immune-associated SNPs show altered accessibility in RO-3306–treated Th1 cells. Top: enhancer–promoter loops; bottom: scATAC-seq signal tracks. For each enhancer, two metrics are displayed: log₂ fold change in accessibility (RO-3306 full/late vs. control, Wilcoxon test; bar color = log₂FC, * = FDR < 0.05) and percentage of cells with non-zero accessibility.

Functional enrichment of the TF regulons highlighted key pathways in T cell biology (**Figure 6B**). CDK1-sensitive TFs IRF8 and STAT1 targeted IFN signaling genes, while BACH2, FOSL2, NR4A2/3, REL, and NFKB1 regulated leukocyte activation and cytokine production, underscoring the regulatory landscape linking signaling to transcription.

To assess how CDK1 activity in CD4+ T-cells is linked to immune-related traits, we tested whether open chromatin regions associated with TF motifs (defined by chromVAR within the SCENIC+ network) were enriched for heritability (stratified linkage disequilibrium score regression, S-LDSC^81^), controlling for all peaks accessible in our scATAC-seq dataset. Peaks linked to the CDK1-sensitive TFs ETS1, IRF8, and LEF1 (**Figure 5G**) were significantly enriched for several traits (**Figure 6C**). ETS1-linked peaks were enriched for “*urinary tract or kidney infection*”, consistent with its role in CD4⁺ T cell homeostasis^74^. IRF8- and LEF1-associated peaks showed enrichment for “*allergy, hypersensitivity or anaphylaxis”* and “*Immunological or systemic disorders”* (defined according to the UK Biobank self-reported illness coding, including allergic traits and systemic autoimmune/connective tissue diseases^82^). Overall, this analysis identified ETS1, IRF8, and LEF1 as CDK1-sensitive regulators that couple kinase signaling to genetic risk.

To dissect how these TFs connect genetic variants in regulatory elements to gene expression, we extracted a subnetwork of enhancer–gene links overlapping SNPs for ‘*immunological or systemic disorder*’ traits (**Figure 6D**). The CDK1-sensitive TF IRF8 was linked to *IFNG* through four SNP-containing enhancers (rs2216163, rs2069727, rs2069716, and rs10878796) (**Figure 6E**). Consistent with reduced IFNG expression under CDK1 inhibition (**Figures 4D and 5F**), accessibility at these enhancers decreased in stimulated Th1 cells treated with RO-3306. Since IRF8 activity is increased upon CDK1 inhibition, this suggests IRF8 may act as a repressive or modulatory factor at *IFNG*. Notably, rs2069727 has been associated with altered *IFNG* expression^83^, underscoring the functional importance of this element. We also identified an IRF8-linked enhancer overlapping rs2239704 at the *TNF* locus (**Figure 6F**). Accessibility was reduced under CDK1 inhibition, mirroring decreased *TNF* expression (**Figure 5F**). rs2239704 has been associated with bronchial asthma and tuberculosis^84^, and non-Hodgkin lymphoma^85^, extending the relevance of CDK1-sensitive circuits to multiple disease contexts.

Together, these results position CDK1 as a signaling hub that bridges phosphorylation with disease-associated regulatory elements, uncovering a kinase-to-genetics axis shaping human T cell function.

## Discussion

CD4⁺ T cell differentiation into diverse Th subsets is governed by signaling cascades, transcriptional programs, and chromatin dynamics. While lineage-defining cytokines and TFs have been characterized, the upstream signaling events that coordinate these responses remain less well defined. Here, we integrated phosphoproteomics, transcriptomics, and epigenomics to systematically map signaling dynamics during human CD4⁺ T cell activation and differentiation.

We uncover a role for post-transcriptional regulation in the earliest phase of activation. After 10 minutes, transcripts enriched for AREs were rapidly downregulated despite stable chromatin accessibility, accompanied by phosphorylation of RBPs. This is consistent with prior reports that resting T cells harbor large pools of untranslated mRNAs, positioning RNA turnover as a key regulator of early activation^86,87^. The absence of this signature upon restimulation suggests that especially naive cells rely heavily on post-transcriptional mechanisms. Among the phospho-RBPs, we confirmed phosphorylation of known regulators of pro-inflammatory transcripts in CD4+ T cells such as ZFP36^36^, and KHSRP^30^, and identified less-characterized candidates including SRRM1. Notably, SRRM1 phosphorylation is inversely regulated by PD-1 signaling^12^ (a pathway known to dampen activation and promote exhaustion), suggesting a role at the interface of activation and inhibitory pathways.

At later stages, we observed coordinated remodeling of chromatin accessibility and transcription, with TF activity involving both canonical regulators (AP-1, NF-κB, STATs) and less-characterized factors (including AP2D, ELK3 and HMGA1 in Th1, and ZEP1 and FOSL1 in iTreg). Several TFs displayed phosphorylation dynamics consistent with their inferred activity, underscoring post-translational control as a key layer in Th subset–specific gene expression. Global phosphoproteomics generally detects relatively few TFs, likely reflecting their low abundance, transient expression, and rapid turnover. However, integrating phosphosite data with chromatin-based TF activity inference provided complementary insights into phosphorylation-dependent regulation. Some well-characterized sites clearly influence TF stability, localization, or DNA binding, but the vast majority remain unannotated^88^, limiting functional interpretation from phosphoproteomics alone. Our multi-omic approach therefore highlights how combining phosphorylation dynamics with chromatin accessibility can reveal regulatory mechanisms otherwise hidden in single-layer analyses.

We identify a role for CDK1, classically defined as a cell-cycle kinase, in effector T cell signaling. Pharmacological inhibition impaired IFN-γ production in Th1 cells, while single-cell analyses revealed selective suppression of pro-inflammatory programs alongside preservation of regulatory signatures. This aligns with prior observations in THP-1 monocytes, where CDKs control IFN-β production by regulating its association with actively translating polysomes^89^. In line with this, our single-cell data showed that CDK1 inhibition altered expression of genes involved in mRNA metabolism, including downregulation of the splicing factor *SRRM1*. These findings suggest that CDK1 shapes T cell activation not only through classical signaling and TF activity, but also via post-transcriptional mechanisms such as RNA translation, splicing and stability. Moreover, rapid proliferation is a hallmark of effective T cell responses, and it may be advantageous for pathways controlling cell cycle to overlap with those governing effector function. Indeed, different antigenic stimuli influence cell cycle dynamics in mouse CD8+ T cells^90^, and effector cytokine expression by differentiating CD4+ cells (including IFN-γ) is partially cell cycle-dependent^91^. These results highlight CDK1 as a dual regulator of cell cycle and effector signaling in human T cells, with potential as a context-dependent therapeutic target.

Our integration of CDK1-regulated TF networks with GWAS data highlights how kinase signaling intersects with genetic risk. Regulatory elements targeted by CDK1-sensitive TFs, including ETS1, IRF8, and LEF1, showed heritability enrichment in immune-related traits, linking kinase-regulated transcriptional networks to human disease susceptibility. Subnetwork analysis identified SNP-containing enhancers at *IFNG* and *TNF* loci whose accessibility was altered under CDK1 inhibition. This extends prior work in systemic lupus erythematosus (SLE), where CDK1 promotes type I IFN signaling and its inhibition dampens IFN-stimulated gene expression^92^. At *IFNG*, we identified an enhancer harboring the autoimmune-associated SNP rs2069727, which displayed reduced accessibility and decreased *IFNG* expression under CDK1 inhibition. Notably, multiple sclerosis (MS) patients carrying the rs2069727*G allele exhibit elevated *IFNG* expression^83^, underscoring the functional relevance of this variant. IRF8 activity increased under CDK1 inhibition and was predicted to bind at this site, suggesting a role as a negative regulator of *IFNG*. A similar pattern was observed at the *TNF* locus, where IRF8 binding overlapped a SNP-containing enhancer with reduced accessibility, consistent with reports that IRF8 can inhibit TNF-ɑ expression^93^. Beyond these loci, our GRN–GWAS integration uncovered additional CDK1-sensitive TF–enhancer–SNP connections, offering a broader resource for dissecting how kinase activity and genetic variation converge on immune traits.

Our study provides one of the most comprehensive multi-omic maps of human CD4⁺ T cell differentiation to date, and the first to integrate global phosphoproteomics with transcriptomic and chromatin profiling in primary human T cells. By linking signaling kinases, TF networks, and genetic variation, we offer a systems-level framework for dissecting T cell biology and a resource for future studies on kinase regulation and immune disease risk.

## Methods

### Human CD4⁺ T Cell Isolation and Culture

Healthy donor buffy coats were obtained from the DRK-Blutspendedienst Baden-Württemberg – Hessen. PBMCs were isolated by density-gradient centrifugation using the Ficoll-Paque Plus solution (GE Healthcare Life Sciences, GE17-1440-02), and stored in liquid nitrogen until T cell isolation. Naive CD4+ T cells were isolated using the EasySep Human Naive CD4+ T cell Isolation kit II (Stemcell Technologies, 17555) with ‘The Big Easy’ EasySep Magnet (Stemcell Technologies, 18001). Naive T cells were either processed directly, or cultured in ImmunoCult-XF T cell Expansion medium (Stemcell Technologies, 10981) in the presence of Human T-Activator CD3/CD28 Dynabeads (Gibco, 11131D, 25µL of Dynabeads per 1x10^6^ T cells), supplemented with Th subset-specific cytokine mixes and 1% Penicillin-Streptomycin (PenStrep, Gibco, 15140122). The following cytokine mixes were used:

- Th0: 10ng/ml IL-2 (Peprotech, 200-02)
- Th1: 10ng/ml IL-2, 50ng/ml IL-12 (Miltenyi Biotec, 130-096-705), 1μg/ml anti-IL-4 (R&D systems, MAB204-100)
- iTreg: 10ng/ml IL-2, 5ng/ml TGF-β1 (Peprotech, 100-21).

After 3 three days of culture, medium was changed to fresh medium supplemented with subset specific cytokines, and cells were split into two new wells. After 5 days, the medium was again changed and the cells were left unstimulated or stimulated with 50ng/ml PMA (Sigma-Aldrich, P8139-1MG) and 1µg/ml ionomycin (Sigma-Aldrich, I0634-1MG) for 4 hours. At five key timepoints (naive, 10 minutes, 5 days, 5 days + 10min, 5 days + 4 hours), samples were collected for bulk RNA-seq, ATAC-seq, phosphoproteomics, SUM-seq and flow cytometry as described below. CD3/CD28 Dynabeads were removed using a magnet prior to downstream processing.

### Bulk RNA-sequencing

For bulk RNA-sequencing, 300.000 cells were collected from each condition and RNA was isolated using the RNeasy Mini Kit (Qiagen, 74104) according to the manufacturer’s protocol. During RNA isolation, on-column digestion of DNA was performed using the RNase-Free DNase Set (Qiagen, 79256). RNA concentration and quality was tested on the 2100 Bioanalyzer system (Agilent), using the RNA 6000 Nano kit (Agilent, NC1783726) according to the manufacturer’s protocol, and 100ng of high quality RNA (RIN scores > 8.8) from each sample was taken for sequencing. Library preparation was performed by the EMBL GeneCore facility. Total RNA was subjected to poly(A) selection using the NEBNext® Poly(A) mRNA Magnetic Isolation Module (New England Biolabs, E7490), followed by cDNA synthesis, end repair, adaptor ligation, and PCR enrichment using the NEBNext® Ultra™ II RNA Library Prep Kit for Illumina® (New England Biolabs, E7770), according to the manufacturer’s instructions. Libraries were indexed and multiplexed, with all 39 samples pooled and sequenced together on a single P3 flow cell using the Illumina NextSeq 2000 platform (150 bp paired-end reads).

### Bulk ATAC-sequencing

Chromatin accessibility profiling was performed using a modified version of the ATAC-seq protocol adapted from Buenrostro *et al*.^94^, using in-house assembled Tn5 transposase^95^. Briefly, 50,000 nuclei were isolated from each sample using a non-ionic detergent-based lysis buffer (0.1% NP-40 (Thermo Scientific, 85124), and 0.01% digitonin (Invitrogen, BN2006)) by incubating on ice for 4 minutes, followed by centrifugation and washing in cold wash buffer (10mM Tris-HCl pH7.5, 10mM NaCl, 3mM MgCl2, 0.10% Tween20, 1% BSA in nuclease free water). Tagmentation was carried out at 37 °C for 30 minutes shaking at 500 rpm, using assembled Tn5 in tagmentation buffer (18.8% DMF (Sigma-Aldrich, 227056), 11.8 mM Mg-acetate, 77.6 mM K-acetate, 38.8mM Tris-acetate, 0.12% NP-40, and protease inhibitor cocktail). Following DNA purification using the MinElute PCR Purification Kit (Qiagen, 28006), libraries were PCR-amplified using NEBNext High-Fidelity 2X PCR Master Mix with custom P5 and indexed P7 primers. To minimize PCR amplification bias, library amplification cycles were optimized using a qPCR side reaction to determine the optimal number of additional cycles prior to saturation (typically 5–9 cycles). Final libraries were size-selected using a 1.2x SPRIselect bead cleanup (Beckman Coulter, B23317) and quantified using a Qubit® 2.0 Fluorometer (Q32866), with the Qubit™ dsDNA HS Assay Kit (Invitrogen, Q32854). Quality and fragment distribution were assessed on the 2100 Bioanalyzer system using the High Sensitivity DNA Kit (Agilent, 5067-4626). Libraries were sequenced on an Illumina NextSeq 2000 using paired-end 75 bp reads.

### Phosphoproteomics Sample Preparation, TMT Labeling, and Mass Spectrometry

Samples were harvested at the indicated time points, and cell pellets were washed twice in ice cold DPBS, snapfrozen on dry ice and stored at -80℃ until processing. Protein samples were subjected to the SP3 protocol^96^. For digestion, trypsin was used in a 1:20 ratio (protease:protein) in 50 mM N-2-hydroxyethylpiperazine-N-2-ethanesulfonic acid (HEPES) supplemented with 5 mM Tris(2-carboxyethyl)phosphine hydrochloride (TCEP) and 20 mM 2-chloroacetamide (CAA). Digestion was carried out overnight at 37°C. Eluted peptides were labeled using TMTpro™ 16plex reagent as previously described^97^. Briefly, 0.5 mg of TMT reagent was dissolved in 45 µL of 100% acetonitrile. Subsequently, 4 µL of this solution was added to each peptide sample, followed by incubation at room temperature for 1 hour. The labeling reaction was quenched by adding 4 µL of a 5% aqueous hydroxylamine solution and incubating for an additional 15 minutes at room temperature. Labeled samples were then combined for multiplexing, desalted using an Oasis® HLB µElution Plate (Waters) according to the manufacturer’s instructions, and dried by vacuum centrifugation. For phosphopeptide enrichment, peptides were taken up in IMAC loading solvent (70% acetonitrile, 0.07% TFA). A small aliquot was used for full proteome analysis. Phosphopeptide enrichment was performed on an UltiMate 3000 RSLC LC system (Dionex) using a ProPac IMAC-10 Column 4 x 50 mm, P/N 063276 (Thermo Fisher Scientific) essentially as described in^98^.

An UltiMate 3000 RSLC nano LC system (Dionex) fitted with a trapping cartridge (µ-Precolumn C18 PepMap 100, 5µm, 300 µm i.d. x 5 mm, 100 Å) and an analytical column (nanoEase™ M/Z HSS T3 column 75 µm x 250 mm C18, 1.8 µm, 100 Å, Waters) was coupled to a Orbitrap Fusion™ Lumos™ Tribrid™ Mass Spectrometer (Thermo). Peptides were concentrated on the trapping column with a constant flow of 0.05% trifluoroacetic acid at 30 µL/min. Subsequently, peptides were eluted via the analytical column using a binary solution system at a constant flow rate of 0.3 µL/min. Solvent A consists of 0.1% formic acid in water with 3% DMSO and solvent B of 0.1% formic acid in acetonitrile with 3% DMSO. The peptides were introduced into the Fusion Lumos via a Pico-Tip Emitter 360 µm OD x 20 µm ID; 10 µm tip (New Objective) and an applied spray voltage of 2.4 kV. The capillary temperature was set at 275°C. Full mass scan was acquired with mass range 375 to1500 m/z in profile mode in the orbitrap with resolution of 120000. The filling time was set at maximum of 50 ms for the full proteome with a limitation of 4x10^5 ions. Data dependent acquisition (DDA) was performed with the resolution of the Orbitrap set to 30000, with a fill time of 110 ms for the phosphoproteome (94 ms for the full proteome) and an AGC target of 200%. A normalized collision energy of 34 was applied. MS2 data was acquired in profile mode. Fixed first was mass was set 110 m/z.

### Single-cell Multi-omic Profiling (SUM-seq)

Simultaneous profiling of transcriptome and chromatin accessibility at single-cell resolution was performed using SUM-seq^6^, with minor adaptations. Two independent SUM-seq experiments (involving two and three donors, respectively) were performed to ensure both technical and biological reproducibility. Naive human CD4⁺ T cells were cultured under Th0, Th1, or iTreg-polarizing conditions in the presence of 2.5µM CDK1 inhibitor (RO-3306) or vehicle control (DMSO). At day 5, cells were either left unstimulated or restimulated for 4 hours with 50ng/ml PMA (Sigma-Aldrich, P8139-1MG) and 1µg/ml ionomycin (Sigma-Aldrich, I0634-1MG). For each condition, ∼0.5 million cells were harvested, and fixed in 3% glyoxal (containing 0.75% acetic acid, pH adjusted to 5.0 using 1 M NaOH) for 7 minutes at room temperature. After fixation, cells were washed in RSB–1% BSA–RNasin buffer (RSB-1% BSA-RI; 10 mM Tris-HCl pH 7.5, 10 mM NaCl, 3 mM MgCl₂, 1 mM DTT, 1% BSA, and 20 μg/mL in-house produced RNasin), then slowly cryopreserved in freezing buffer (50 mM Tris-HCl pH 7.5, 5 mM Mg-acetate, 0.1 mM EDTA, 25% glycerol) using a Mr. Frosty™ Freezing Container (Thermo Scientific, 5100-0001), and stored at –70 °C until processing.

For further processing, cryopreserved cells were thawed slowly on ice and washed with RSB–1% BSA–RI buffer. Nuclei were extracted by resuspending cells in lysis buffer (10mM Tris–HCl pH 7.5, 10mM NaCl, 3mM MgCl2, 0.1 % Tween20, 20µg/mL RNasin, 1mM DTT, 1% BSA, 0.025% NP-40 and 0.01% Digitonin), incubated for 1 minute on ice, and washed with wash buffer (10mM Tris–HCl pH 7.5, 10mM NaCl, 3mM MgCl2, 0.1% Tween20, 20µg/mL RNasin, 1mM DTT and 1% BSA). Nuclei counts from a subset of samples were quantified using 4,6-diamidino-2-phenylindole (DAPI) and tryphan blue staining, and downstream workflow was performed with approximately 40,000 nuclei per sample.

Custom barcoded Tn5 transposomes were assembled in-house as described by Lobato-Moreno et al^8^, and Tn5 transposase was produced following an in-house protocol^95^. Fixed nuclei were subjected to in situ transposition using custom barcoded Tn5 transposomes, followed by reverse transcription with barcoded oligo(dT) primers. 12% PEG 8000 (v/v; Jena Bioscience, CSS-256) was included in the RT mix. Following the RT reaction, all samples were pooled in RSB-RI and washed twice with RSB–1% BSA-RI, counted, and 700.000 nuclei were taken for cDNA/mRNA hybrid tagmentation. After gap filling and exonuclease digestion, nuclei were filtered through a 40µM Flowmi cell strainer (Sigma-Aldrich, BAH136800040), and counted. 350.000 nuclei were loaded onto the Chromium Controller (10x Genomics) according to the 10x Genomics single-cell ATAC standard protocol using the Chromium Next GEM Single Cell ATAC Kit v2 (10x Genomics, 1000390), in a mixture containing barcoding reagent B, reducing reagent B, barcoding enzyme, and 500nM of a blocking oligonucleotide. Samples were loaded on the Chromium Next GEM Chip H (10x Genomics,1000162) following the standard 10x workflow. Barcoded emulsions were recovered, cleaned, and subjected to pre-amplification and modality-specific final library amplification. Library concentrations were measured using Qubit 1× dsDNA-HS (Thermo Scientific, Q33230) and fragment distributions were assessed using the Bioanalyzer High Sensitivity DNA kit (Agilent, 5067-4626). Sample specific barcodes are provided in **Supplemental Table 1**.

Final scATAC and scRNA libraries were sequenced on an AVITI Cloudbreak High platform (Element Biosciences) using paired-end 150 bp reads (ATAC: read 1, 55 cycles; index 1, 11 cycles; index 2, 16 cycles; read 2, 55 cycles; RNA: read 1, 95 cycles; index 1, 6 cycles; index 2, 16 cycles; read 2, 21 cycles).

### Flow Cytometry

For intracellular cytokine detection, cells were restimulated with PMA (50 ng/mL) and ionomycin (1µg/mL) or left unstimulated, in the presence of Brefeldin A (1:000, Biolegend, 420601). For proliferation assays, naive CD4⁺ T cells were labeled prior to culture with 5 μM CellTrace™ Blue Cell Proliferation Kit (Invitrogen, C34574) according to the manufacturer’s instructions. Cells were harvested at indicated time points, washed with ice cold DPBS, and stained for viability using Zombie Aqua Fixable Viability Dye (BioLegend, 423101) for 10 minutes at room temperature in the dark. For intracellular staining, cells were fixed and permeabilized using the FOXP3/Transcription Factor Staining Buffer Set (Invitrogen, 00-5523-00), for 45 minutes at 4°C, followed by staining with Pacific Blue™ anti-human IFN-γ Antibody (dilution 1:50, BioLegend, 502521) Alexa Fluor® 647 anti-T-bet Antibody (dilution 1:100, Biolegend, 644803), and PE/Dazzle™ 594 anti-human FOXP3 Antibody (dilution 1:50, Biolegend, 320125) for 30 minutes at 4°C. Samples were acquired on the BD FACSymphony™ A3 (BD Biosciences), and analyzed using FlowJo v10.9.

### Bulk RNA-sequencing data analysis

Raw RNA-sequencing data was processed with our in-house RNA-Seq data processing Snakemake pipeline as previously described^39^. Briefly, raw reads were first assessed with FastQC (http://www.bioinformatics.babraham.ac.uk/projects/fastqc/) and trimmed using cutadapt^99^. Trimmed reads were aligned to the GRCh38/hg38 reference genome using STAR^100^, and gene-level expression was quantified using both featureCounts^101^ (mapQ ≥ 10, no multi-overlap counting), and expression values were summarized as raw counts. Raw gene-level count matrices were processed in R using DESeq2^102^ with the design formula “∼ replicate + condition”, controlling for biological replicate variation. Genes with fewer than 10 total counts across all samples were filtered out. Variance-stabilizing transformation (VST) was applied to normalized counts for PCA, identifying top varying genes. Differential expression analysis was performed using the Wald test, as implemented in DESeq2, and genes with log₂ fold change > 0.58 or < 0.58 and adjusted p-value < 0.05 were considered differentially expressed.

### Bulk ATAC-sequencing data analysis

Raw ATAC-sequencing data was processed with our in-house ATAC-Seq data processing Snakemake pipeline as previously described^39^. Briefly, raw reads were first assessed with FastQC (http://www.bioinformatics.babraham.ac.uk/projects/fastqc/), trimmed using Trimmomatic^103^ with MINLEN cutoff of 20, and aligned to the GRCh38/hg38 reference genome using Bowtie2^104^ (“--very-sensitive”, maximum fragment length 2000 bp). Only properly paired reads with MAPQ ≥10 were retained using SAMtools^105^. PCR duplicates were identified and removed using Picard’s MarkDuplicates (http://broadinstitute.github.io/picard). Reads containing indels or soft-clipping were excluded by filtering the CIGAR string (CIGAR: “ID”) to retain only exact matches. Peaks were called using MACS2^106^ with the parameters “--nomodel --shift -100 --extsize 200 -q 0.01”, and read counts per peak were quantified using featureCounts^101^. A consensus peak set was generated using DiffBind^107^, considering peaks detected in at least three samples (minOverlap = 3). Raw read counts were quantified across the consensus peaks using dba.count() with score = DBA_SCORE_READS and without re-centering peaks on summits (summits = FALSE), and filtered to retain autosomal peaks only. Raw consensus peak counts were first upper-quartile normalized using the EDASeq package^108^, with normalization factors calculated from the betweenLaneNormalization() function using “which = “full”, and “offset = TRUE”. These normalization factors were then applied to a DESeqDataSet object via normalizationFactors(). Differential accessibility in DESeq2 was modeled using a design formula accounting for biological replicate and condition (“∼ Replicate + Condition”), and peaks with log₂ fold change > 0.58 or < 0.58 and adjusted p-value < 0.05 were considered differentially accessible. Peak annotation was performed using ChIPseeker^109^ with the TxDb.Hsapiens.UCSC.hg38.knownGene database.

### Phosphoproteomics data analysis

Raw spectral data were processed using FragPipe with MSFragger, searching against the UniProt Homo sapiens proteome with appended common contaminants and reversed sequences for FDR estimation. Fixed modifications included carbamidomethylation on cysteine and TMT labeling on lysine residues; variable modifications included phosphorylation on serine, threonine, and tyrosine (STY), oxidation of methionine, and N-terminal acetylation or TMT labeling. Trypsin was specified as the protease, allowing up to two missed cleavages. The precursor and fragment mass error tolerance was set to 20 ppm. False discovery rate (FDR) control was set at 1%.

The raw output files of FragPipe^110^ were processed using the R programming language (ISBN 3-900051-07-0). Only peptide spectral matches (PSMs) with a phosphorylation probability greater 0.75 and proteins with at least 2 unique peptides were considered for the analysis. Phosphorylated amino acids were marked with a * in the amino acid sequences behind the phosphorylated amino acid, labeled with a 1, 2 or 3 for the number of phosphorylation sites in the peptide and concatenated with the protein ID in order to create a unique ID for each phosphopeptide. Raw TMT reporter ion intensities were summed for all PSMs with the same phosphopeptide ID. Transformed summed TMT reporter ion intensities were first cleaned for batch effects using the ‘removeBatchEffects’ function of the limma package^111^ and further normalized using the vsn package (variance stabilization normalization)^112^. Missing values were imputed with ‘knn’ method using the Msnbase package^113^. The replicate information was added as a factor in the design matrix given as an argument to the ‘lmFit’ function of limma. Also, imputed values were given a weight of 0.05 in the ‘lmFit’ function. Phosphopeptides were considered significantly regulated if they exhibited an FDR below 0.05.

### Gene Set Enrichment Analysis

Gene ontology (GO) enrichment analysis was performed using clusterProfiler^114^ to identify biological processes enriched among differentially expressed or accessible genes, or differentially phosphorylated proteins. For each contrast, genes with an adjusted p-value < 0.05 were selected, and enrichment was carried out using the enrichGO() function with the org.Hs.eg.db annotation database and keyType = “SYMBOL”. The background universe consisted of all tested genes for that contrast. To reduce redundancy among GO terms, results were simplified using the simplify() function with a similarity cutoff of 0.7 based on adjusted p-values.

### Differential TF Activity Analysis (diffTF)

Differential TF activity was assessed using diffTF^39^ with default parameters, using motifs from the HOCOMOCOv12 (invivo) database. As input, we used consensus ATAC-seq peaks identified via DiffBind (as described above), together with the corresponding ATAC-seq BAM files. The analysis was run in analytical mode, using DESeq2 for statistical testing. TFs were considered significantly differentially active at an FDR < 0.05.

### Co-phosphorylation network analysis using WGCNA

To identify coordinated phosphorylation modules across T cell subsets and timepoints, we applied Weighted Gene Co-expression Network Analysis (WGCNA)^65^ to phosphoproteomics data. Normalized phosphosite intensities across all samples were used, retaining only phosphosites significantly regulated (FDR < 0.05) in any differential analysis condition. A signed co-expression network was constructed using a soft-thresholding power of 8. The topological overlap matrix (TOM) was computed and hierarchical clustering was performed to group phosphosites based on their co-regulation patterns. Modules were identified using dynamic tree cutting with a minimum module size of 30 and merged using a module eigengene correlation threshold of 0.75 (cut height = 0.25). Module eigengenes were calculated to summarize the expression profiles of each module across samples. These were used to assess expression patterns across T cell subsets (Th0, Th1, iTreg) and stimulation conditions over time. Phosphosite-level Z-scores were visualized per module and per cell type to summarize co-phosphorylation dynamics. In addition, modules were annotated with module membership (kME) scores to highlight hub-like phosphosites contributing to module structure.

### Kinase Set Enrichment Analysis (KSEA)

To infer kinase activities from phosphoproteomics data, we performed Kinase Set Enrichment Analysis (KSEA) using the KSEAapp package^60^. Input data consisted of differentially phosphorylated peptides derived from limma analyses. For each peptide, the following attributes were provided: UniProt accession (Protein), gene symbol (Gene), peptide identifier (Peptide), phosphosite(s) (Residue.Both, e.g., “S102”), adjusted p-value (p), and fold change (FC, non-log-transformed). KSEA scores were computed for each comparison using the KSEA.Scores() function with NetworKIN integration (NetworKIN=TRUE, NetworKIN.cutoff=5). Kinase scores were calculated for both the normalized full phosphoproteome and phospho-only datasets. Z-scores were retained only for kinases with at least three substrate phosphosites (m > 2), and values with adjusted p-value > 0.05 were set to zero to highlight only significant activity changes.

### AU-rich element (ARE) enrichment analysis

Gene-level annotations of AREs were obtained from the AREsite2^20^ database, including both 3′UTR- and intron-localized elements. DE gene lists from each RNA-seq comparison were overlapped with AREsite2 annotations to flag ARE-containing transcripts. For each comparison, we tested whether AREs were significantly enriched among downregulated DEGs (adjusted p-value < 0.05 and log₂ fold change < 0) using Fisher’s exact test, comparing the frequency of AREs in DEGs versus non-DEGs.

### SUM-seq data analysis

Raw sequencing data were processed using the SUM-seq analysis pipeline (https://git.embl.de/grp-zaugg/sum-seq_analysis).

For the RNA modality, base calls were converted to FASTQ format using bcl2fastq. The cell barcode (i5) was concatenated to the sample index and UMI (Read 2), followed by demultiplexing using Je^115^, allowing for up to two mismatches in the index. Reads were aligned to the GRCh38/hg38 reference genome using STARsolo^116^ in paired-end and unstranded mode. The CB_UMI_Simple protocol was used to extract cell barcodes (CB) and UMIs (UB), and UMIs were deduplicated using the 1MM_All method. Gene expression was quantified using exonic reads and Gencode v34 annotations. Gene expression matrices generated by STARsolo were imported using the ReadSTARsolo function from the Seurat package^117^, and all sample-level Seurat objects were merged into a single object using Merge_Seurat_List from the scCustomize package. Quality control filtering was applied to retain cells if they expressed between 200 and 10,000 genes, had ≤15% mitochondrial gene expression, and >500 total RNA counts. Two pre-filtered Seurat objects representing independent experiments were first normalized using log-normalization (LogNormalize, scale factor = 10,000), followed by identification of the top 5,000 variable features (vst method), scaling with regression of mitochondrial gene content (percent.mt), and PCA. Integration anchors were then computed using the first 50 principal components, and datasets were integrated with IntegrateData.

For the ATAC modality, base calls were converted to fastq format and demultiplexed by i7, allowing for one mismatch, using bcl-convert. Demultiplexed reads were aligned to the hg38 genome and fragment file generation was performed with chromap^118^, and fragment files were processed using ArchR^119^. A merged ArchR project was created by combining .arrow files from the two independent experiments. High-quality single cells were retained by applying filtering criteria of at least 1,000 fragments, TSS enrichment score ≥ 5, and promoter ratio ≥ 0.1. Dimensionality reduction was performed on the filtered ArchR project using the addIterativeLSI() function with two iterations, 25,000 variable features, and dimensions 2–30. To correct for potential batch effects between the independent experiments, Harmony integration was applied via addHarmony() using the reduced dimensions from LSI and grouping by donor identity. Pseudo-bulk replicates were generated by aggregating cells by condition using addGroupCoverages(), and peak calling was performed with MACS2 via addReproduciblePeakSet(), using a reproducibility cutoff of 3, up to 100,000 peaks per group, and a q-value threshold of 0.05. The resulting consensus peak set was used to construct a cell-by-peak count matrix with addPeakMatrix(). To assess peak signal quality, the fraction of reads in peaks (FRiP) was computed per cell, and cells with FRiP<0.3 were excluded from further analysis.

To integrate single-cell RNA and ATAC modalities, processed ArchR and Seurat objects were subset to include only shared barcodes. ATAC data were extracted as a ChromatinAssay using the ArchR peak matrix and peak set, and merged into the Seurat object alongside RNA data. RNA counts were SCTransform-normalized with donor and experiment regressed out, followed by PCA and Harmony integration. ATAC data were normalized using TF-IDF and reduced with SVD, with batch correction applied using Harmony. Weighted nearest neighbors (WNN) integration was performed on the RNA and ATAC Harmony embeddings, and multimodal UMAPs were generated to visualize integrated cell states across conditions and donors. Differential gene expression analysis was performed using the Seurat function FindMarkers, with the MAST test to account for zero-inflated single-cell data. Donor and experiment were included as latent variables to control for batch effects. Differential chromatin accessibility was assessed using a Wilcoxon rank-sum test implemented in the presto R package. Each comparison was tested independently, genes and peaks with a false discovery rate (FDR) below 0.05 were considered differential.

### TF Motif Activity Inference (ChromVAR)

To quantify TF motif activity, the chromVAR algorithm^120^ was applied via ArchR using the previously generated fragment files and peak calls. ChromVAR deviations were computed using a motif position frequency matrix from the HOCOMOCOv12 database^121^, and the resulting per-cell deviations and z-scores were extracted from the ArchR project. These were integrated into the multimodal Seurat object as separate assays (chromVAR and chromVAR_Z). Differential motif activity between conditions was assessed using the FindMarkers function in Seurat, applied to the chromVAR_Z assay with a logistic regression (LR) test and experiment and donor as latent variables. Motifs with FDR < 0.05 were considered significantly different.

### SCENIC+ Analysis

SCENIC+ analysis^80^ was based on the integrated Seurat object containing both RNA and ATAC modalities, and was run in multiome mode using an in-house Snakemake pipeline. The ATAC data was processed using the cisTopic module with 40 topics to identify patterns of chromatin accessibility across cells. Differentially accessible regions were identified using a log2 fold-change threshold of 0.5 and adjusted p-value < 0.05. Motif enrichment analysis was performed with cisTarget, using the v10nr_clust_public motif database and including both direct and orthology-based annotations. Enriched motifs were filtered using similarity FDR < 0.001, a minimum of 3 motif hits, and a minimum AUC threshold of 0.005 for motif activity inference. Enhancer-to-gene and TF-to-gene links were inferred using gradient boosting (GBM) based importance scores, and Spearman correlation for region–gene associations. Finally, enhancer gene regulatory networks (eGRNs) were assembled and filtered based on multiple quantile thresholds and minimal region/gene link constraints.

### Cell Cycle Scoring

Cell cycle phase distributions were assessed from single-cell counts using the CellCycleScoring() function in Seurat, stratified by treatment, stimulation condition. For each Th subset, the number of cells in G1, S, and G2/M phases under control-treated, stimulated and unstimulated conditions were calculated. Proportions were derived by normalizing cell counts within each group to the total number of cells per condition. To test for differences in the distribution of cells across cell cycle phases between stimulated and unstimulated conditions, we used a two-sample test for equality of proportions with continuity correction (prop.test() function in R). The test compared the proportion of cells in each phase between conditions, using total cell counts as denominators. P-values were corrected for multiple comparisons using the Benjamini–Hochberg false discovery rate method.

### Linkage Disequilibrium Score Regression

Linkage Disequilibrium Score Regression (LDSC) was used to test for enrichment of SNP heritability in chromatin accessibility peaks associated with specific TFs. Specifically, we evaluated enrichment in peaks associated with IRF8, LEF1, and ETS1, as identified by chromVAR analysis, using all T cell peaks considered for chromVAR as the background set. Heritability enrichment was tested across 8,890 traits from the 22.10 release of Open Targets Genetics^122^, including GWAS from the UK Biobank, FinnGen, and other repositories. We applied Stratified LDSC (S-LDSC)^81^ to traits with estimated SNP heritability (h²) greater than 0.04 and a case count exceeding 10,000. Traits were filtered for those annotated as related to immune system disease, hematologic disease, or infectious disease. Heritability enrichment was calculated using default settings, and p-values were corrected for the number of peak sets tested per trait using the Benjamini–Hochberg false discovery rate method.

### Overlap of TF-regulated peaks with GWAS SNPs and network visualization

BED files corresponding to chromVAR peaks associated with IRF8, ETS1, and LEF1 were imported and converted to GRanges objects. Summary statistics for GWAS traits were obtained from the Open Targets Genetics and harmonized to include rsIDs. SNP genomic positions were retrieved using the Ensembl BioMart interface (GRCh38), and converted into GRanges objects. We then used findOverlaps() to identify SNPs intersecting TF-specific peaks. To link chromVAR-inferred TFs to putative regulatory targets, we intersected the GWAS-overlapping peaks with TF–target links inferred from SCENIC+. Peaks were parsed from the SCENIC+ GRN table and matched to nearby SNPs. For each TF–gene pair, we computed the number of distinct enhancers (peaks) and the total number of overlapping GWAS SNPs. These statistics were used to generate TF–gene networks with edge weights reflecting SNP burden and enhancer support. Networks were constructed using the igraph^123^ package and visualized with ggraph in R. For selected TF–gene pairs, we visualized the genomic loci using plotgardener^124^.

## Data availability

(Single cell) RNA and ATAC-sequencing data will be made available at the European Genome and Phenome Archive, and phopshoproteomics data will be made available at the proteomics identifications (PRIDE) database upon publication.

## Code availability

Code is deposited at https://github.com/NilaServaas/Tcell_Differentiation_2025, and will be made public upon publication.

## Acknowledgements

We thank the EMBL Genomics Core Facility for help with sequencing, the Protein Purification Core Facility for Tn5 and RNasin production, the Flow Cytometry Core Facility for assistance with FACS analysis, the Proteomics Facility for help with phosphoproteomics experiments, and the EMBL IT team for access to the HPC cluster. We also thank members of the Zaugg group at EMBL for their input and scientific discussions, and K.D. Prummel and S. Lobato-Moreno for their support with ATAC-seq and SUM-seq experiments. N.H.S. and J.B.Z. acknowledge funding from GSK through the EMBL–GSK collaboration framework (3000032294), and from EMBL’s Infection Biology Transversal Theme. This project is co-funded by the European Union (ERC, EpiNicheAML, 101044873) to J.B.Z. Views and opinions expressed are however those of the authors only and do not necessarily reflect those of the European Union or the ERC. Neither the European Union nor the granting authority can be held responsible for them.

## Contributions

N.H.S., M.F.S., and J.B.Z. conceived and designed the study. N.H.S. performed all experiments, with assistance from H.G.B. for single-cell experiments. L.A. and N.H.S. collected peripheral blood mononuclear cells from healthy donors. J.J.S. processed samples for phosphoproteomic analysis. N.H.S. carried out all bioinformatic analyses. A.C. contributed to the S-LDSC analysis, I.B. to the SCENIC+ analysis, and F.S. to the phosphoproteomics analysis. N.H.S. and J.B.Z. wrote the manuscript with input from J.P.R. J.B.Z. supervised the project with input from M.F.S., L.E., and J.P.R. All authors reviewed and approved the final manuscript.

## Ethics declarations

Human biological samples were sourced ethically, and their research use was in accord with the terms of the informed consents under an IRB/REC approved protocol.

---

**Supplemental Figure 1.**
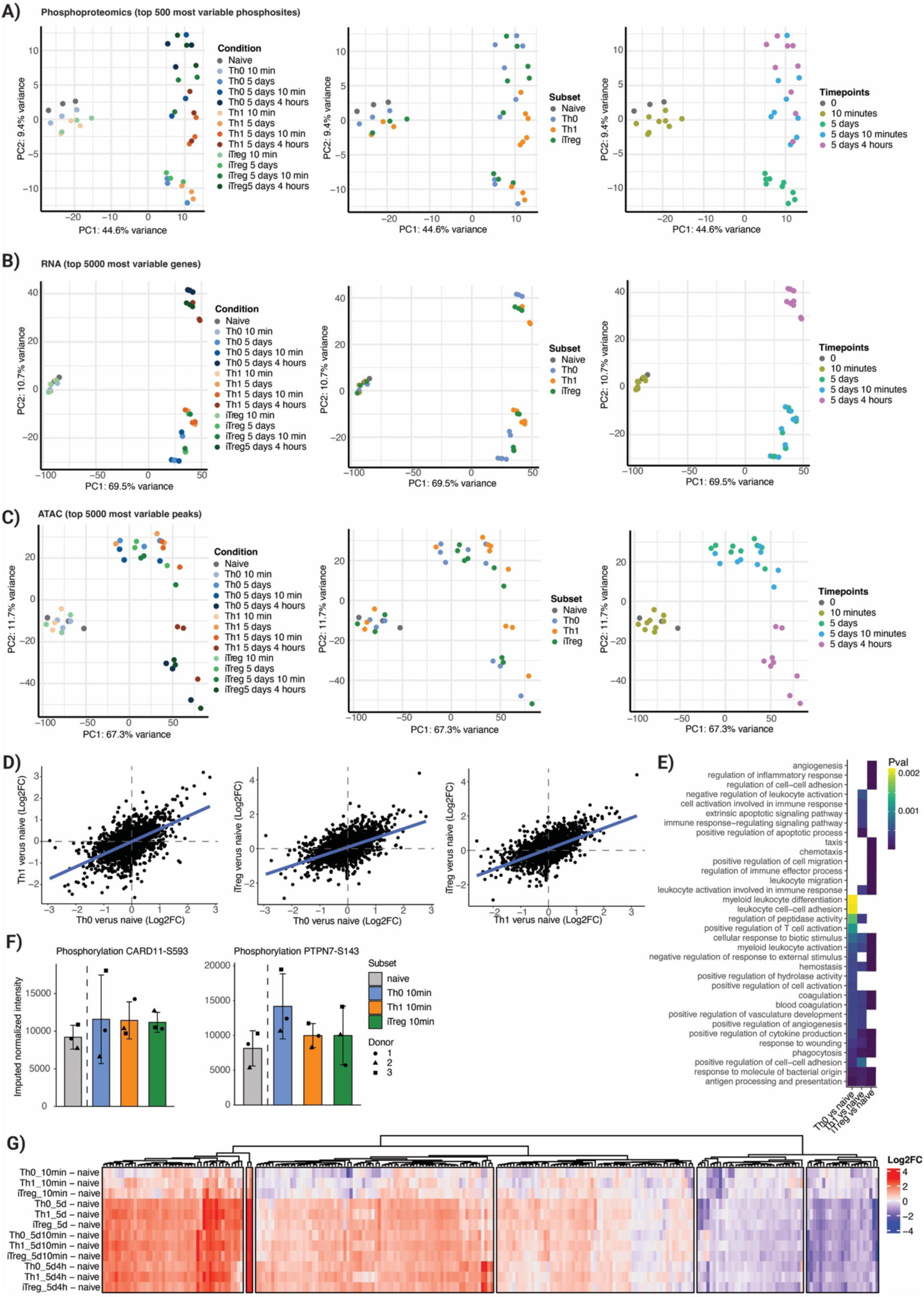
Global profiling of phosphorylation, gene expression, and chromatin accessibility across T cell activation and differentiation. **(A–C)** Principal component analysis (PCA) of the top 500 most variable features across all conditions for (**A**) phosphoproteomics, (**B**) RNA-seq, and (**C**) ATAC-seq. Each point represents one sample. Left panels show samples colored by condition, middle by Th subset, and right by stimulation timepoint. **(D)** Correlation of fold changes (obtained from limma) of differentially phosphorylayed sites in Th0, Th1 and iTregs after 10 minutes of initial stimulation compared to naive T cells. **(E)** Gene ontology enrichment analysis of differentially expressed genes 10 minutes after initial activation compared to naive T cells. Top 10 terms (based on P-value) for each Th subset are shown. **(F)** Barplots showing phosphorylation dynamics of CARD11 (S593) and PTPN7 (S143), which are upregulated within 10 minutes of stimulation. Bars show the mean ± standard deviation of normalized phosphosite intensities across donors, with individual replicates indicated by shape. **(G)** Heatmap showing global phosphorylation changes of differential RNA bindings proteins (RBPs) detected across all Th subsets and timepoints (colors indicate log2 fold-change of pairwise comparisons versus naive). Rows represent phosphosites and are clustered by similarity in log2 fold-change patterns.

**Supplemental Figure 2.**
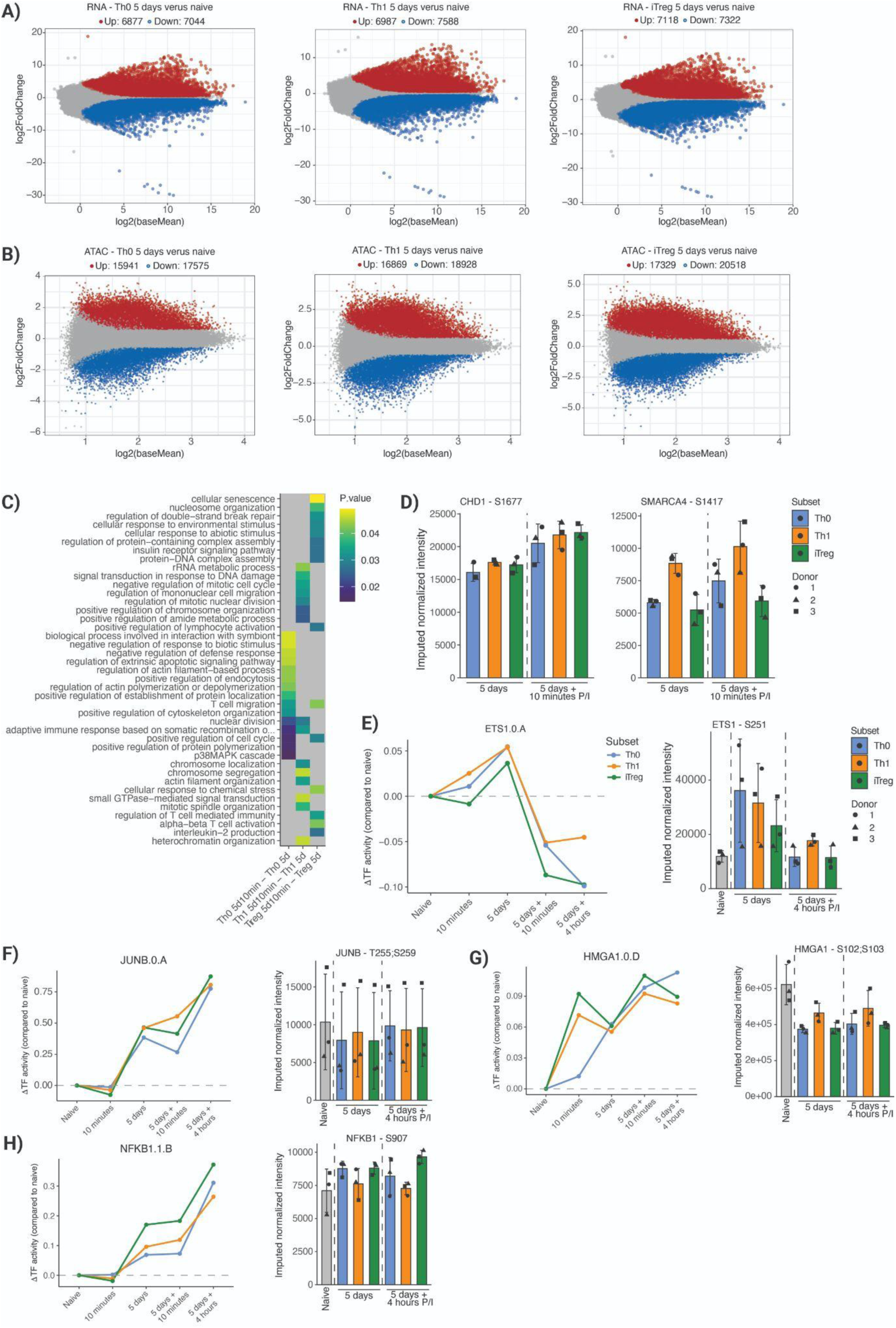
Multi-omics profiling reveals widespread transcriptional and chromatin remodeling during T cell differentiation and phosphorylation-associated regulation of transcription factor (TF) activity. **(A)** MA plots showing differential gene expression (RNA-seq) between naive CD4⁺ T cells and cells cultured for 5 days under Th0, Th1, or iTreg-polarizing conditions. Significantly upregulated (red) and downregulated (blue) genes are highlighted (adjusted p-value < 0.05, log2FC -0.58< or >0.58). **(B)** Differential chromatin accessibility (ATAC-seq) in the same comparisons as in (A). **(C)** Gene ontology enrichment analysis of differentially phosphorylated proteins 10 minutes after PMA/ionomycin restimulation. **(D)** Barplots showing normalized phosphorylation intensities of chromatin-associated proteins CHD1 (S1677) and SMARCA4 (S1417) across subsets at 5 days and 5 days and 10 minutes post-restimulation (P/I = PMA/ionomycin). Data from three donors are shown. **(E–H)** TF activity (from diffTF) and phosphosite intensity measurements for selected TFs across conditions. Left: diffTF-inferred TF activity over time, compared to naive. Right: phosphorylation levels of key regulatory sites in ETS1 (S251), JUNB (T255/S259), HMGA1 (S102/S103), and NFKB1 (S907).

**Supplemental Figure 3.**
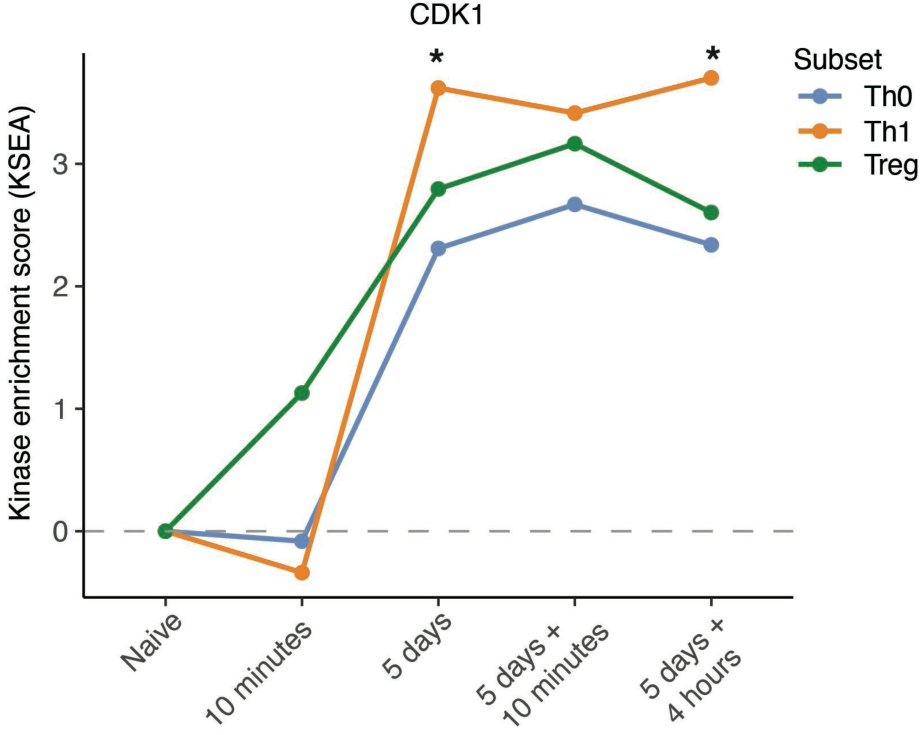
KSEA-inferred CDK1 activity across naive, differentiated, and restimulated CD4⁺ T cell subsets. CDK1 kinase enrichment scores were calculated based on phosphoproteomics data. Asterisks indicate timepoints within which Th1 cells have a significant enrichment score over Th0 and iTreg (NetworKIN (cutoff ≥ 5), adjusted p-value < 0.05 and minimum of 3 substrates).

**Supplemental Figure 4.**
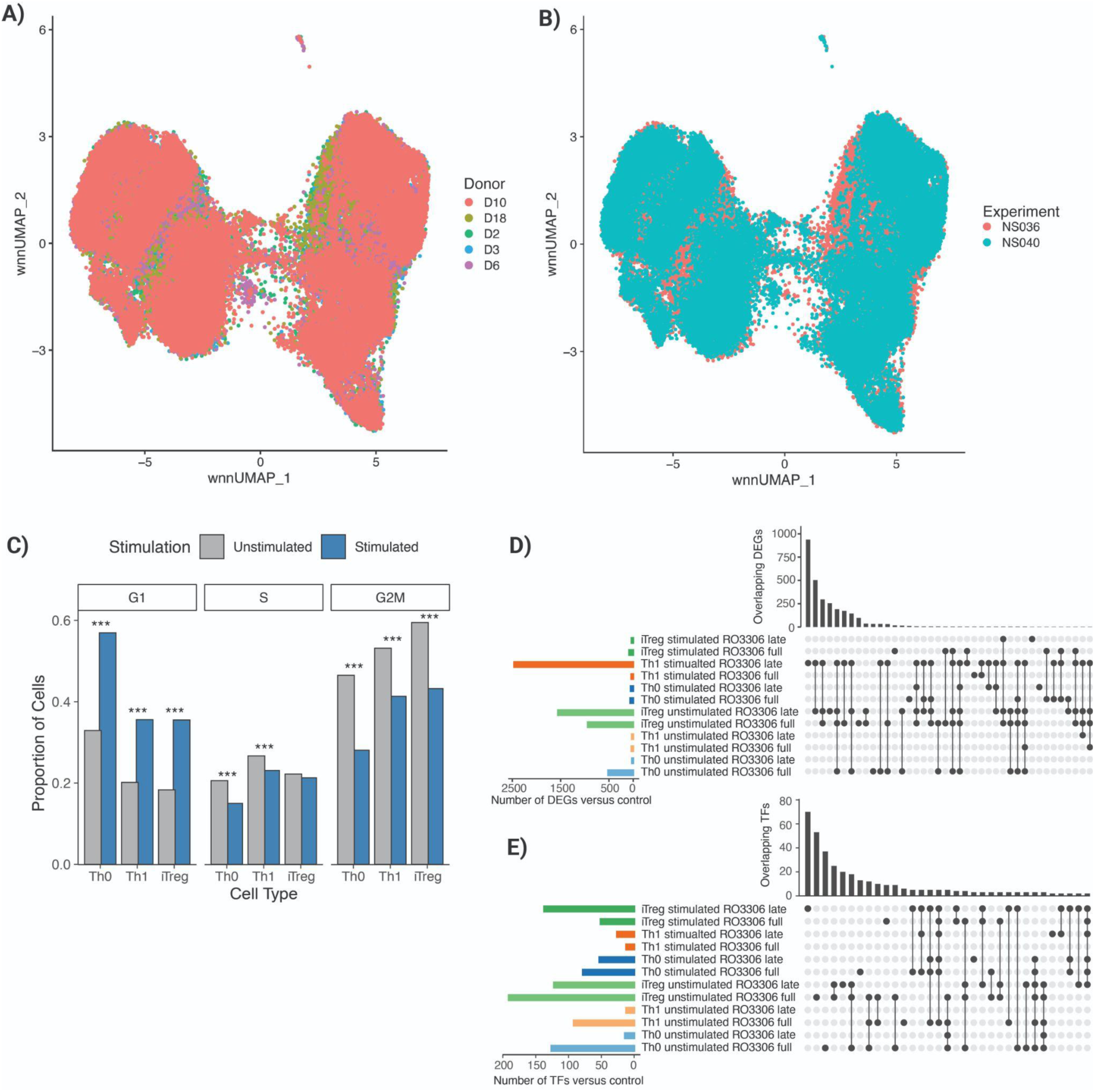
CDK1 inhibition induces transcriptional and chromatin remodeling across donors and experiments. **(A)** Weighted nearest neighbours (WNN) UMAP of integrated scRNA-seq and scATAC-seq data showing cells colored by donor and **(B)** experimental replicates, confirming biological and technical reproducibility. **(C)** Bar plots showing proportions of cells in each cell cycle phase (G1, S, G2/M) for stimulated and unstimulated cells across Th0, Th1, and iTreg subsets (***p < 0.001, two-sample test for equality of proportions with continuity correction with Bonferroni correction). **(D)** UpSet plot displaying the number and overlap of differentially expressed genes (DEGs) in RO3306– treated versus control cells across all conditions. **(E)** UpSet plot of differentially active transcription factors (TFs, chromVAR) in RO3306–treated versus control cells across all conditions.

**Supplemental Table 1.**
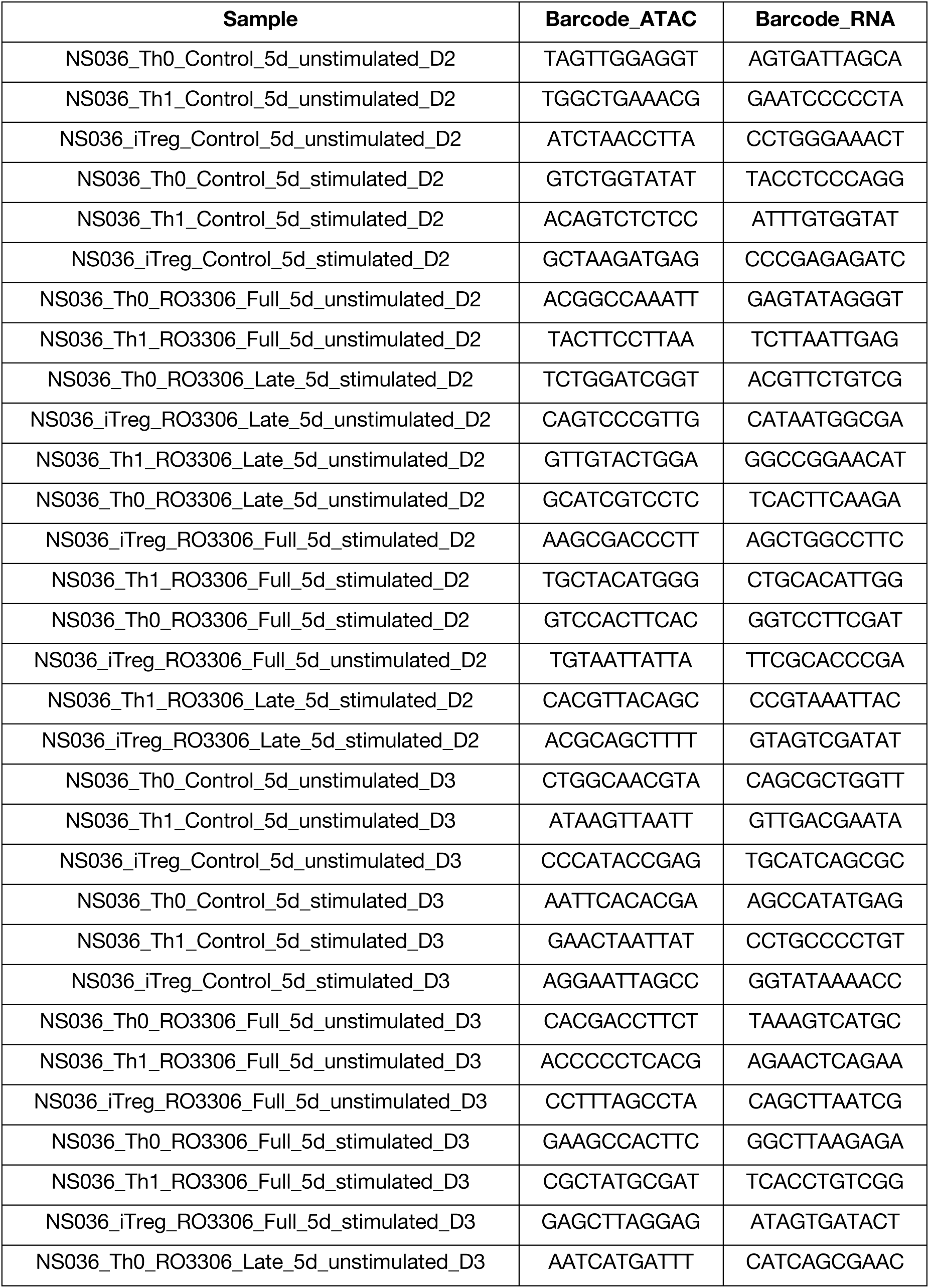
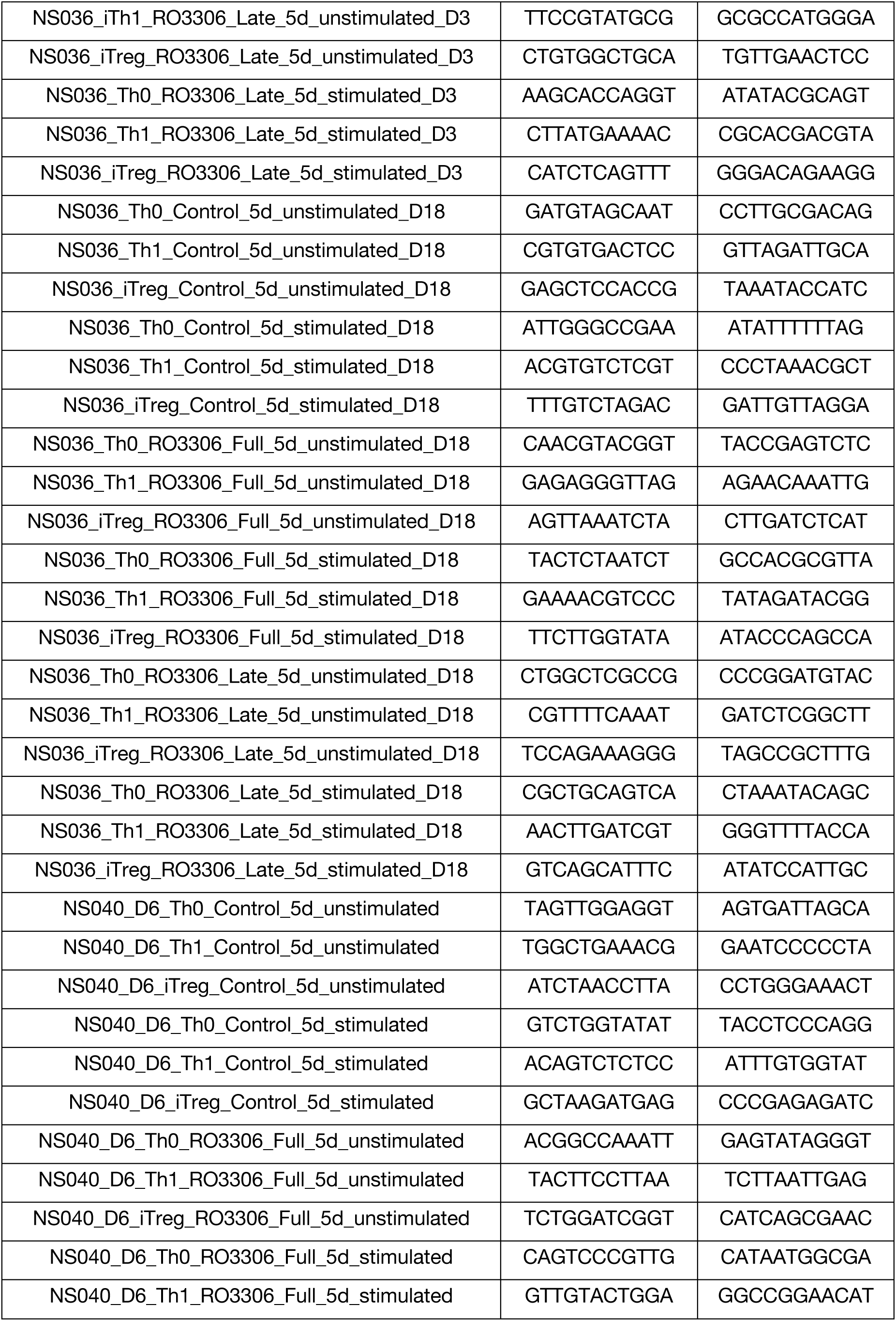
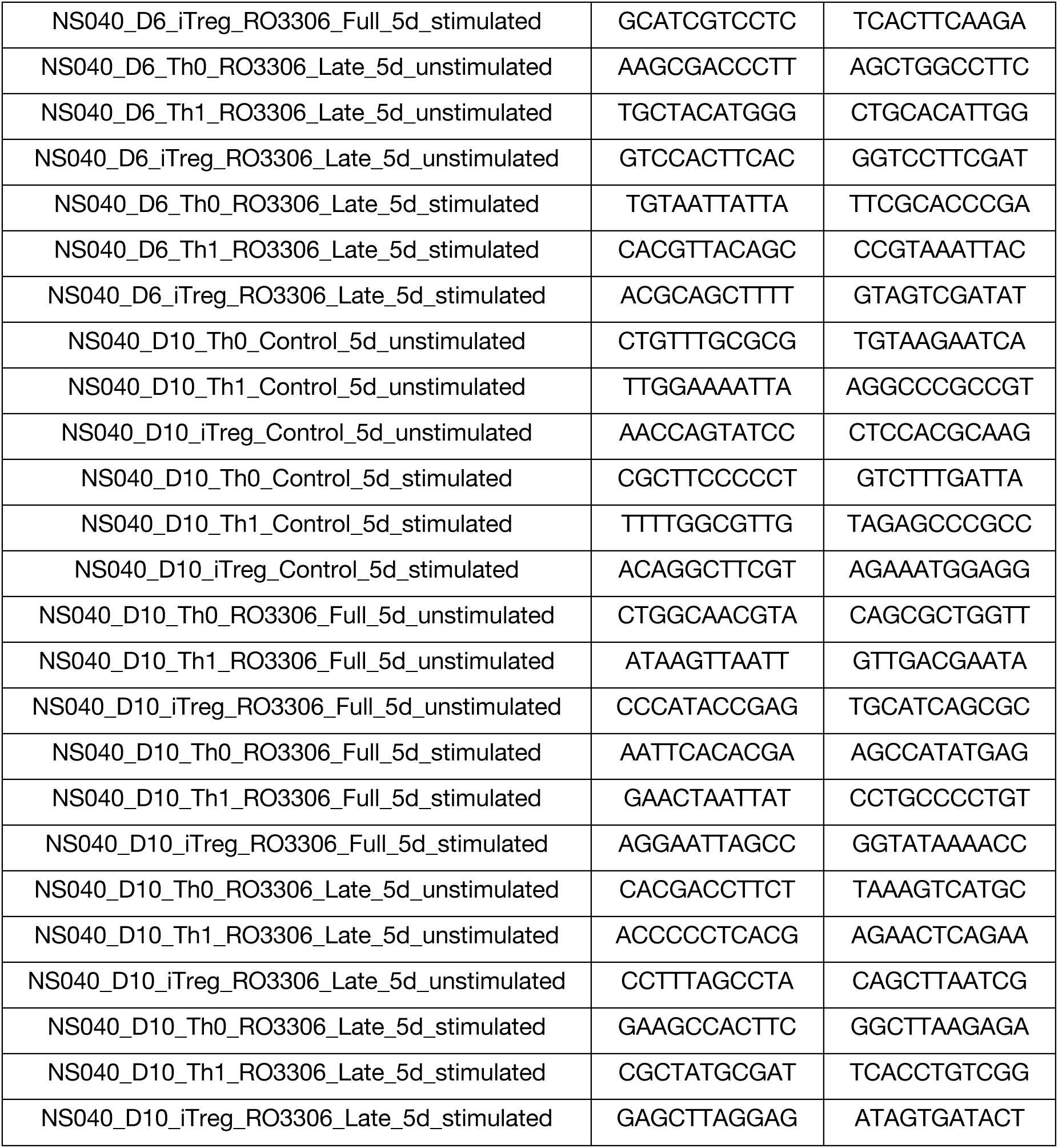
Sample specific barcodes for demultiplexing of SUMseq data.

