## Supplementary figures and images for "Integrative Multi-Omics Identifies CDK1 as a Key Signaling Regulator of CD4^+^ T Cell Effector Function"

### Supplemental Figure 1

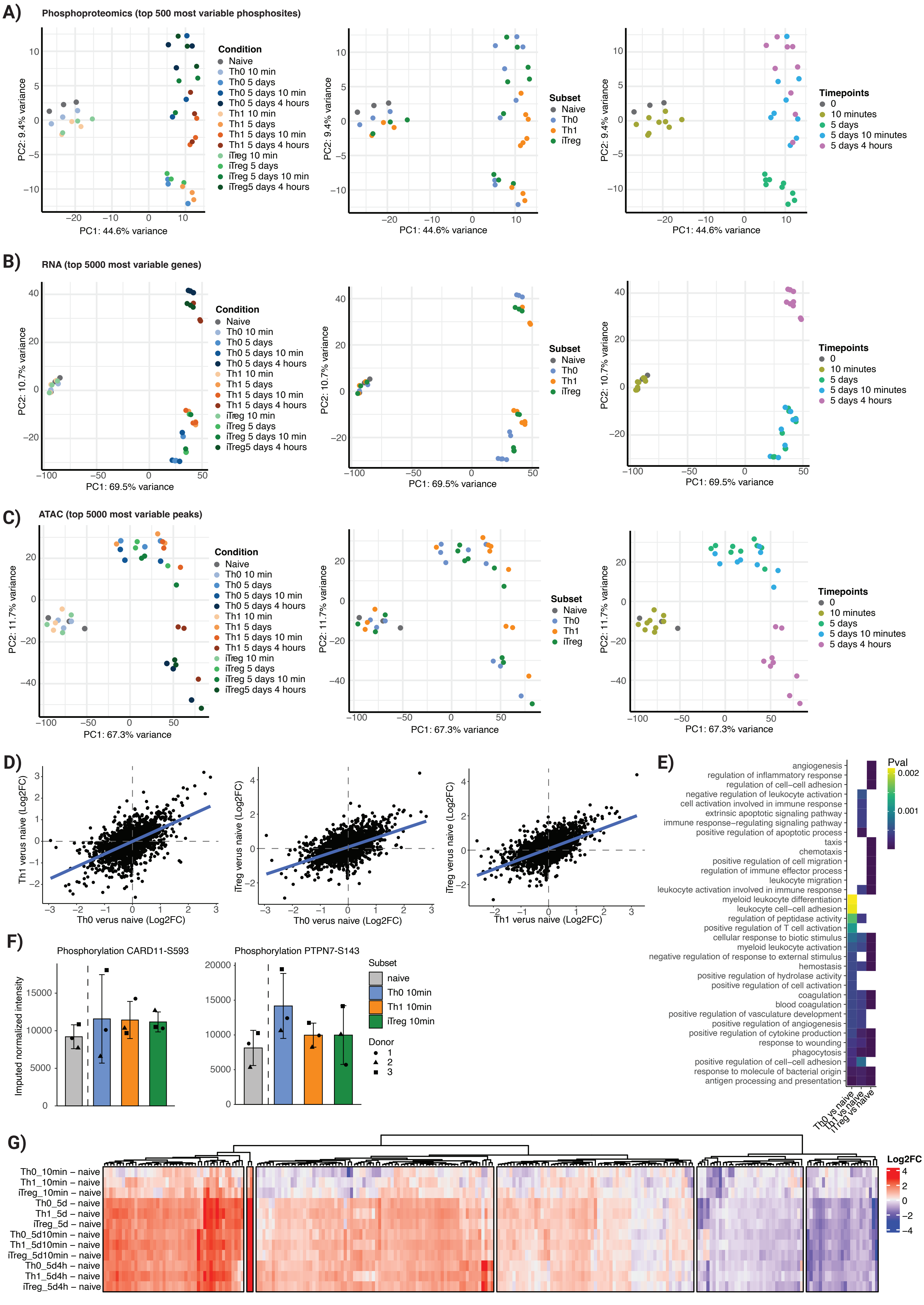

### Supplemental Figure 2

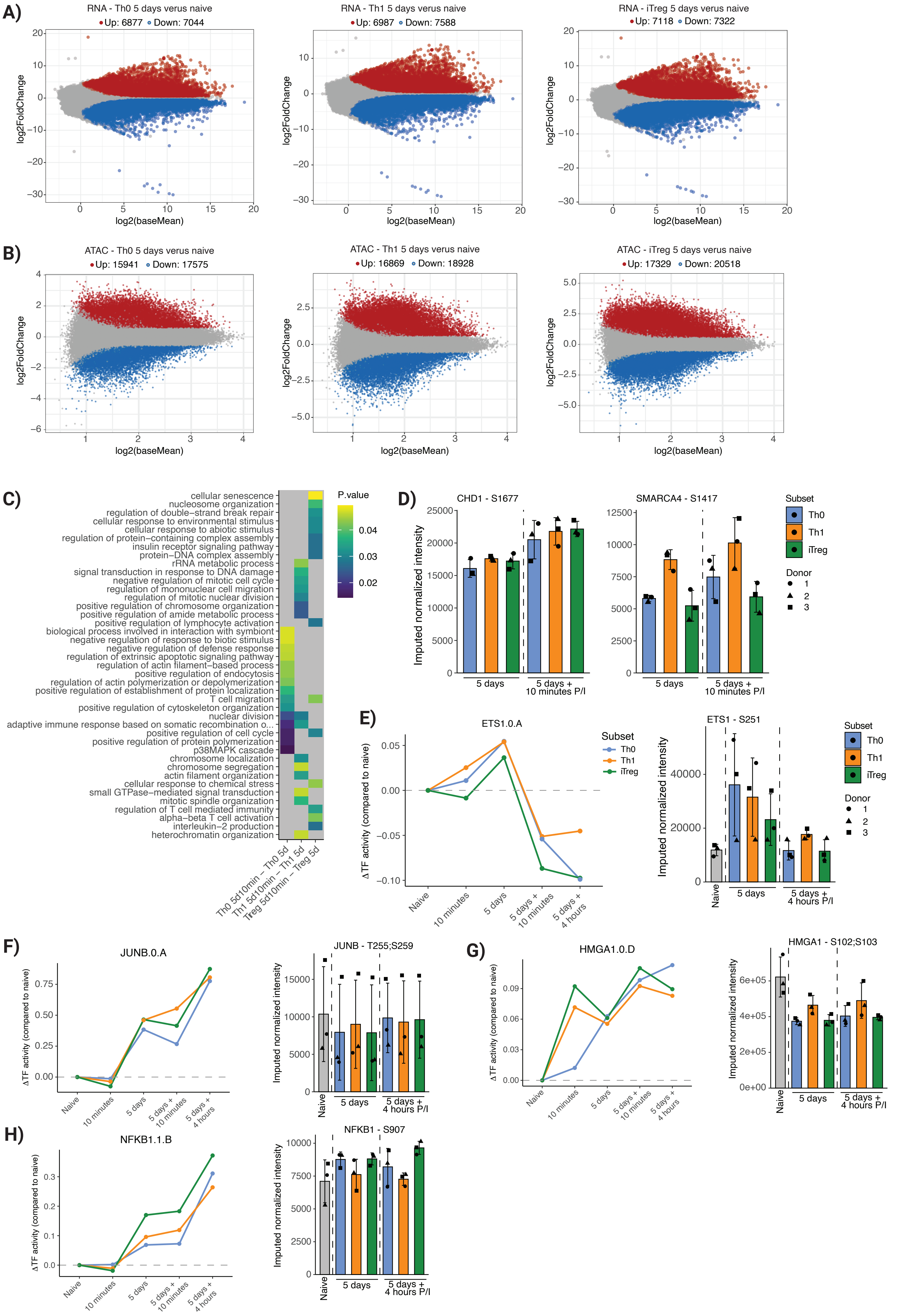

### Supplemental Figure 3

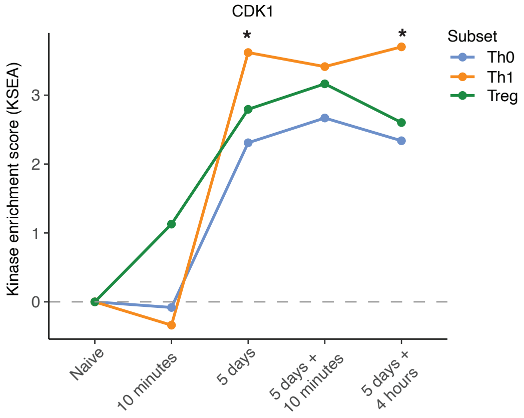

### Supplemental Figure 4

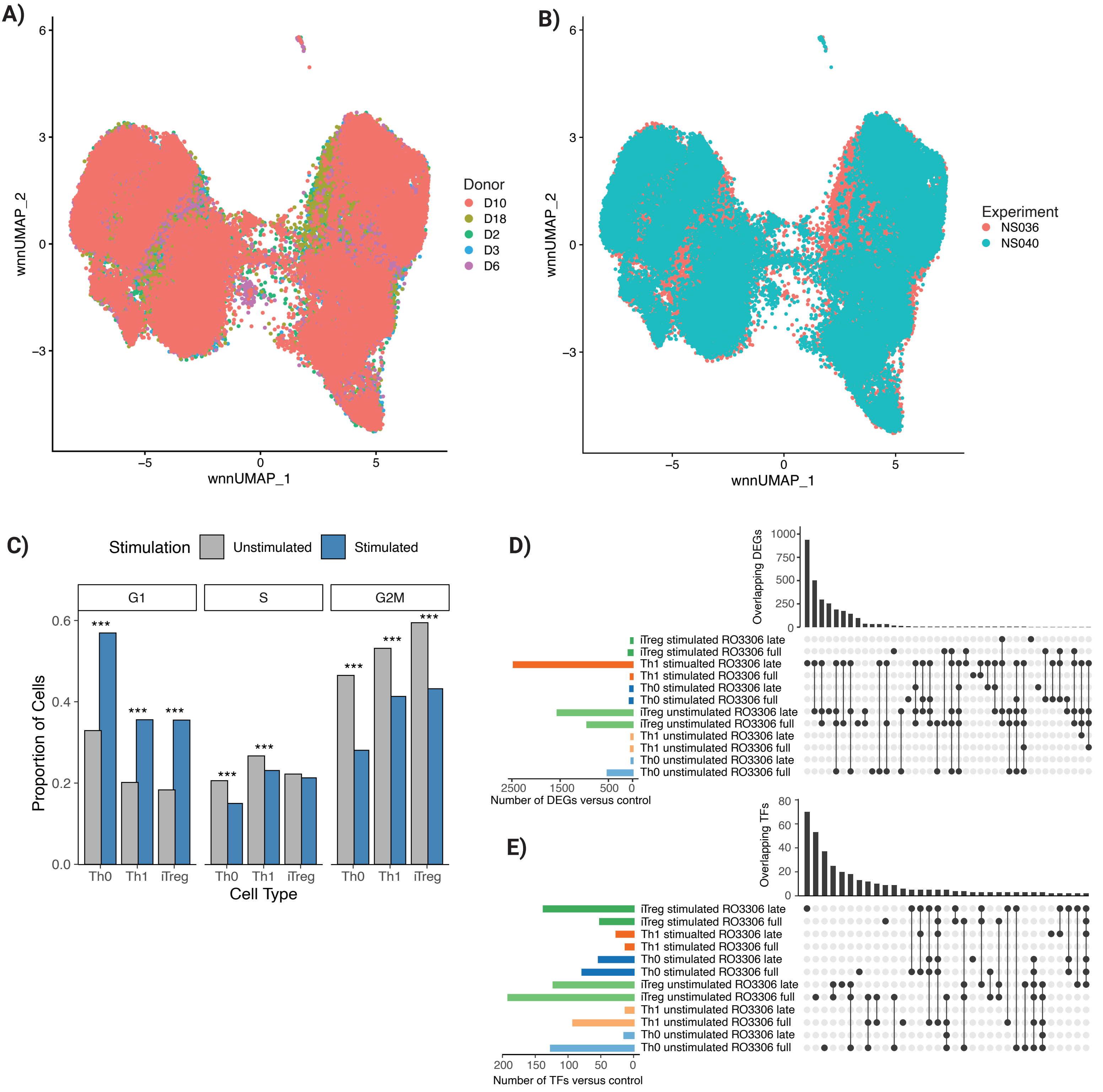
